# Association of Whole-Person Eigen-Polygenic Risk Scores with Alzheimer’s Disease

**DOI:** 10.1101/2022.09.13.507735

**Authors:** Amin Kharaghani, Earvin Tio, Milos Milic, David A. Bennett, Philip L. De Jager, Julie A. Schneider, Lei Sun, Daniel Felsky

**Affiliations:** Krembil Centre for Neuroinformatics, Centre for Addiction and Mental Health, Toronto, ON, Canada; Division of Biostatistics, Dalla Lana School of Public Health, University of Toronto, Toronto, ON, Canada; Institute of Medical Science, University of Toronto, Toronto, ON, Canada; Rush Alzheimer’s Disease Center, Rush University Medical Center, Chicago, IL, USA; Centre for Translational and Computational Neuroimmunology, Columbia University Medical Center, New York, NY, USA; Department of Statistical Sciences, University of Toronto, Toronto, ON, Canada; Department of Psychiatry, University of Toronto, Toronto, ON, Canada

**Keywords:** Polygenic risk scores, Alzheimer’s disease, WGCNA, eigen-PRS, whole-person

## Abstract

Late-Onset Alzheimer’s Disease (LOAD) is a heterogeneous neurodegenerative disorder with complex etiology and high heritability. Its multifactorial risk profile and large portions of unexplained heritability suggest the involvement of yet unidentified genetic risk factors. Here we describe the “whole person” genetic risk landscape of polygenic risk scores for 2,218 traits in 2,044 elderly individuals and test if novel eigen-PRSs derived from clustered subnetworks of single-trait PRSs can improve prediction of LOAD diagnosis, rates of cognitive decline, and canonical LOAD neuropathology. Principal component analyses of thousands of PRSs found generally poor global correlation among traits. However, component loadings confirmed covariance of clinically and biologically related traits and diagnoses, with the top PCs representing autoimmune traits, cardiovascular traits, and general pain medication prescriptions, depending on the PRS variant inclusion threshold. Network analyses revealed distinct clusters of PRSs with clinical and biological interpretability. Novel eigen-PRSs (ePRS) derived from these clusters were significantly associated with LOAD-related phenotypes and improved predictive model performance over the state-of-the-art LOAD PRS alone. Notably, an ePRS representing clusters of traits related to cholesterol levels was able to improve variance explained in a model of brain-wide beta-amyloid burden by 1.7% (likelihood ratio test *p*=9.02×10^−7^). While many associations of ePRS with LOAD phenotypes were eliminated by the removal of *APOE*-proximal loci, some modules (e.g. retinal defects, acidosis, colon health, ischaemic heart disease) showed associations at an unadjusted type I error rate. Our approach reveals new relationships between genetic risk for vascular, inflammatory, and other age-related traits and offers improvements over the existing single-trait PRS approach to capturing heritable risk for cognitive decline and beta-amyloid accumulation. Our results are catalogued for the scientific community, to aid in the generation of new hypotheses based on our maps of clustered PRSs and associations with LOAD-related phenotypes.

## Introduction

Late-Onset Alzheimer’s disease (LOAD) is the most common cause of dementia in older adults [1]. It is a highly polygenic and multifactorial disease [2] with heritability estimated to be between 58% to 79% [3]. Despite the identification of its canonical pathologies decades ago, no effective disease-modifying treatments are available. This is likely a result of the etiological complexity of the illness and challenges associated with studying causality for a cascade of events that precede the emergence of symptoms by decades. High-throughput genetic approaches and the ongoing era of genome-wide association study (GWAS) discovery have provided clues into the complex, potentially causal mechanisms of LOAD [4] and highlighted disturbances across multiple processes and systems. The most recent genetic evidence points toward inflammatory, neurotrophic, metabolic, and cellular senescence mechanisms [5–9], and genetic correlations have been observed between LOAD and a multitude of physical, psychiatric, and other neurological illnesses [8,10].

From an epidemiological perspective, factors affecting risk for and progression of LOAD are highly varied. A systematic review [11] on LOAD risk factors highlighted statin use, alcohol consumption, diet, educational attainment, physical and cognitive activity, head injury (in males only), age, diabetes, estrogen therapy, smoking, and social engagement as major risk factors (in addition to genetic risk and protective factors, such as *APOE* ε4 and ε2, respectively). A recent review [12] of factors specifically affecting the progression of LOAD identified six with cohesive evidence (malnutrition, genetic variants, altered gene regulation, baseline cognitive level, neuropsychiatric symptoms, and extrapyramidal signs), though nine more showed conflicting associations in the literature. Additional factors with less clear, potentially bidirectional, associations with LOAD include a large host of often comorbid conditions including sleep disturbances, depression, and gastrointestinal, cardiovascular, and other inflammatory diseases [13,14]. Adding to the complexity of contributing risk factors, the age at which some emerge can affect an individual’s LOAD risk profile. For example, obesity [15], hypertension [16], high cholesterol [17] and statin use [18] in mid-life are associated with an increased risk of dementia later in life. Importantly, these lifestyle, environmental, clinical, and behavioural factors are themselves heritable, and exert their neuropathological effects via multiple biochemical cascades which may be unique to each person [19]. This calls for a more sophisticated multivariate, multi-trait analyses of the genetic component of risk for predicting LOAD, incorporating the whole-person scope of potential phenotypic contributions.

To our knowledge, there have only been a small handful of studies attempting to unravel the phenome-wide genetic risk landscape of LOAD using polygenic approaches. One group used results from the UK Biobank PheWAS [20] to identify patterns of correlated phenotypes and individual genetic variants across thousands of human traits, including LOAD, using singular value decomposition applied to summary statistics [21]. Similar work, focusing specifically on LOAD, proposed another summary statistics-focused method [22], identifying 48 PRSs each with significant association with LOAD genetic risk. Other analyses from the Alzheimer’s Disease Genomics Consortium (ADGC) have examined multifactorial risk for LOAD by calculating PRSs for 22 candidate modifiable risk factors, finding only an educational attainment PRS significantly associated with LOAD [23]. Here, we build on previous efforts in four important ways: 1) by performing dimensionality reduction on phenome-wide PRSs calculated on a population of elderly, rather than operating on summary statistics alone, 2) by examining how broad patterns of phenotypic overlap across PRSs are affected by the adjustment of SNP inclusion thresholds (including sensitivity analyses with the inclusion and exclusion of the *APOE* and the MHC genetic regions), 3) by deriving and testing novel composite eigen-PRSs (ePRSs) through the application of the weighted gene correlation network analysis (WGCNA), and 4) by evaluating the added value of these novel ePRSs explicitly against the standard LOAD-specific PRS [24] for association with longitudinal cognitive decline and post-mortem neuropathological burden.

We hypothesize that a set of phenome-wide PRSs will yield discrete intra-correlated clusters of related traits and that patterns of correlation and clustering will change as a function of SNP inclusion criteria, possibly reflecting between-trait differences in polygenicity. We also hypothesize that the proposed composite ePRSs, derived from the WGCNA algorithm, will capture genetic risk from proximal, possibly overlapping etiological risk factors for LOAD thus improving the prediction of LOAD and LOAD-related phenotypes over the single-PRS method.

## Materials and Methods

### The Religious Orders Study (ROS) and Memory Aging Project (MAP) Participants

ROS and MAP are two longitudinal cohort studies based out of the Rush Alzheimer’s Disease Centre (RADC) in Chicago (IL) [25] designed with harmonized protocols to be analyzed together. Participants are older, recruited free of known dementia, and agree to annual detailed clinical evaluation, cognitive testing, and organ donation at the time of death. A total of 2,044 participants were determined to be of European ancestry and included in our analyses (see **Supplementary Methods**). All phenotypic data are available via controlled access through the RADC Resource Sharing Hub (www.radc.rush.edu) and genomics data are available through Synapse (https://www.synapse.org/#!Synapse:syn3219045). The Institutional Review Board of Rush University Medical Center approved each study, and all subjects provided informed, written consent and signed an Anatomical Gift Act. All participants signed a repository consent for resource sharing.

### Clinical diagnosis of LOAD

For all subjects, available clinical data were reviewed at the time of death by a neurologist with expertise in dementia. Summary diagnoses were made blinded to all post-mortem data and case conferences with one or more neurologists and a neuropsychologist were used to achieve consensus when required. Diagnoses fell into one of six possible categories: no cognitive impairment (NCI), mild cognitive impairment (MCI) with or without secondary cause of impairment, probable late onset Alzheimer’s disease (LOAD; NINCDS criteria [26]), possible LOAD (with additional cause of impairment), and other primary cause of dementia [27]. For the purposes of our case-control modelling of consensus diagnosis at time of death, we considered only probable LOAD and NCI participants; all other analyses of neuropathological and cognitive outcomes included all participants with available data.

### Assessment of longitudinal cognitive decline

At each annual evaluation, all subjects were administered 21 cognitive tests of which 19 were used in both ROS and MAP [28]. The raw scores from these 19 tests were standardized to z-scores across the cohort and averaged to yield the global cognitive function summary score at each study visit. Slopes of change in global cognitive performance over time (cognitive decline) were calculated per subject using linear mixed modelling, with a random effect for study visit, controlling for age at baseline, as previously described [29].

### Assessment of postmortem beta-amyloid and tau neuropathology

Board-certified neuropathologists examined all brains without prior knowledge of clinical or demographic information. Brain removal and fixing procedures have been detailed extensively [30]. Complete data from neuropathological examinations were available for up to 1,252 participants at time of study. Immunohistochemistry was used to quantify total beta-amyloid and paired helical filament tau (PHFtau) across eight brain regions (hippocampus, entorhinal cortex, midfrontal cortex, inferior temporal, angular gyrus, calcarine cortex, cingulate cortex, frontal cortex), as described previously [31]. The square root transformed brain-wide averages of beta-amyloid and tau pathologies were used as in previous analyses of these data [32]. **Figure S1** illustrates the distribution of the pathological measurements alongside global cognitive decline of the ROS/MAP participants.

### Genotyping and imputation

Genotype data were available for 2,067 ROS/MAP participants, between two batches: n_batch1_=1,686 genotyped using the Affymetrix GeneChip 6.0 and n_batch2_=381 genotyped using the Illumina OmniQuad Express platform; batch was included as a covariate in our analyses. Details of raw genotype quality control (QC) have been previously described [33]. Briefly, prior to submission for imputation, genotypes were preprocessed using the TOPMed Imputation Server-recommended data preparation pipeline (https://topmedimpute.readthedocs.io/en/latest/prepare-your-data.html). Each batch was imputed separately using the TOPMed Imputation Server (TOPMed reference r^2^)[34], including Eagle (v2.4) for allelic phasing and Minimac4 (v1.5.7) for imputation. Imputed output data from the TOPMed server for each batch were filtered for imputation quality (removing SNPs with r^2^<0.8) before merging and mapping to rsIDs (dbSNP build 155). This resulted in a set of 9,329,439 high-quality, bi-allelic autosomal SNPs. Additionally, we verified the European ancestral background of genotyped participants by mapping the multidimensional scaling (MDS) plots of the ROS/MAP SNP genotypes against the 1000 Genome project reference populations (phase 3) [35] (details in **Supplementary Methods** and **Figure S2**). Prior to our PRS calculation, additional sample and variant quality controls were performed, removing SNPs and subjects according to the following criteria:

- SNPs with minor allele frequency (MAF) < 0.01 (1,549,210 variants)
- SNPs with Hardy-Weinberg Equilibrium (HWE) *p*-value < 1×10^−6^ (132 variants)
- SNPs with missingness rate > 0.01 (978 variants)
- Individuals with no cognitive or neuropathological phenotype data available were removed (7 individuals)
- Individuals with higher-than-expected rates of SNP heterozygosity within 3 standard deviations of the mean (15 individuals)
- Individuals with non-white European ancestry were removed (1 individual) Following QC, a total of 2,044 individuals and 7,723,915 variants remained for analysis.

### Selection of GWAS summary statistics from the UK Biobank

For phenome-wide PRS calculations, we used GWAS summary statistics derived from the UK Biobank, a large-scale population-based study including approximately 500,000 individuals [36]. The Pan-UK Biobank Consortium (https://pan.ukbb.broadinstitute.org/) has recently conducted GWAS for all available study phenotypes with sufficient statistical power. Out of a total of 7,221 phenotypes with GWAS performed in the European ancestry sample (which matches our sample of genetically verified European-ancestry ROS/MAP participants), we selected 2,218 phenotypes that had heritability estimates greater than or equal to 5% and were sex non-specific. The heritability scale of each phenotype was provided by Pan-UKBB consortium (calculated using SAIGE [37]), and selected sex non-specific phenotypes using their provided meta-data (“pheno_sex” column indicating “both”). Additional manual selection was performed to remove phenotypes with no cases for either sex (indicated by columns “n_cases_full_cohort_females” or “n_cases_full_cohort_males”), indicating geographical or administrative information, or duplicate indications or codings of the same underlying measurement. The full list of selected phenotypes for our study are provided in **Table S1**, and further details of selection criteria are outlined in **Supplementary Methods**. For calculating a benchmark PRS for LOAD, we also downloaded summary statistics from the latest LOAD GWAS, published by Bellenguez et al. [38]

### Calculation and compilation of whole-person PRS matrices

PRSice (v2.3.3) [39] was used to generate PRSs across all traits at 15 different SNP inclusion *p*-value thresholds (α-values) including 1 (all SNPs), 0.1, 0.05, 0.01, 5 × 10^−3^, 1 × 10^−3^, 5 × 10^−4^, 1 × 10^−4^, 5 × 10^−5^, 1 × 10^−5^, 5 × 10^−6^, 1 × 10^−6^, 5 × 10^−7^, 1 × 10^−7^, and 5 × 10^−8^ (only genome-wide significant SNPs).

For each α-value threshold, we obtained a set of PRSs for up to 2,218 traits (fewer traits are included at more stringent thresholds, as they may not have identified any SNPs reaching that threshold). For PRS calculations, we used the default “average” method of PRSice [40]. To account for patterns of linkage disequilibrium (LD), we applied a 500Kb sliding window and r^2^ threshold of 0.2 during variant clumping, as in previous work [41]. To control for fine population structure [42], the first ten principal components of genotype data and a variable representing genotype batch were regressed out of each score independently. We then conducted further QC, removing any PRS with fewer than five contributing SNPs. We also identified any pairs of PRSs with extremely highly correlation (Pearson correlation > 0.95) and retained only one (phenotypes with ICD10 code were preferred otherwise selected randomly). Finally, each PRS was standardized (mean subtracted and then divided by standard deviation) to obtain z-scores.

This resulted in 15 sets (for the 15 different α-value thresholds) of PRS “matrices” for downstream analysis. Each PRS “matrix” is a *n* × *k* matrix of phenome-wide PRSs calculated across traits, where *n* = 2,044 is the number of participants after QC, and *k* ≤ 2,218 is the number of traits. The value of *k* varies across the 15 sets (outlined in **Table S2**), because at a given α-value threshold, some PRSs may be constructed from fewer than five SNPs thus removed as part of the QC.

Notably, previous work has observed a profound influence of variants in the *APOE* region of chromosome 19 on LOAD phenotypes in PRS analyses [43]. Additionally, many psychiatric and neurological disorders are strongly associated with genetic variants in the major histocompatibility complex (MHC) region of chromosome 6 [44], which is also notoriously challenging to impute due to complex LD [45]. To assess the influence of these regions on the clustering of PRSs and resulting associations with LOAD-related phenotypes, we additionally calculated all the 15 sets of PRSs excluding a) the MHC region (GRCh37: chr6:25477797-36448354) [21] (55,204 variants), and b) the *APOE* region (GRCh37: chr19:45300000-45500000) [46] (622 variants).

This resulted in a total of 15 × 4 = 60 sets of phenome-wide PRS matrices, for the 15 different α-value thresholds and four different *APOE* and/or *MHC* inclusion and exclusion criteria: **1)** including all genotyped and imputed variants genome-wide (primary analysis; denoted “ALL”), **2)** removing only *APOE* SNPs (denoted “APOE-”), **3)** removing only MHC SNPs (denoted “MHC-”), and **4)** removing both *APOE* and MHC SNPs (denoted “MHC/APOE-”). For clarity, we denote these PRS matrices as *Π*_*subscript*_, where the *subscript* indicates whether the matrix was calculated on the whole genome or with indicated regions excluded (i.e. ALL, MHC-, APOE-, MHC/APOE-) and the *α*-value threshold used for SNP inclusion (e.g. 1 × 10^−4^, 5 × 10^−8^). For example, 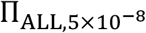 is for the matrix when a PRS was calculated without SNPs in the MHC region and at an α-value threshold of 1 × 10^−8^. **Table S2** outlines the dimension of each PRS matrix after QC.

### Principal component analysis (PCA) of whole-person PRS matrices

PCA is a multivariate dimension-deduction technique that can describe the covariance among a set of variables (in our case, PRSs) [47]. All PCA analyses were performed using R (v4.2.0) (https://www.r-project.org/). PCA was first applied to each of the 60 PRS matrices separately to identify the major variance components of phenome-wide genetic risk and describe the contributors to each top principal component. This exploratory PCA analysis served three purposes: 1) as a cursory test of accuracy for the large sets of PRS calculations and post-processing (do PCs load onto known epidemiologically and biologically related traits?), 2) to inform expectations for cohesion in downstream clustering analyses, and 3) to provide confirmation of observations made previously based on GWAS summary statistics that the majority of phenome-wide variance in genetic risk for human traits is explained by genetic risk for metabolic and cardiovascular health-related measures [21].

### Clustering of PRS matrices and calculation of eigen-PRSs (ePRS) using WGCNA

To identify meaningful, discrete clusters of correlated PRSs across the phenome for ePRS construction, we used the weighted gene co-expression network analysis (WGCNA) framework [48],which leverages the network properties of biological phenomena to cluster input measures. Although WGCNA was originally developed for use in gene expression experiments to identify biologically coherent co-expressed gene modules, the principles on which it operates are not unique to such measures; WGCNA has been used to build epigenetic networks [49] and to subtype patient populations [50]. Notably, the original authors of WGCNA have also adapted the method for network-based analyses of GWAS summary statistics (the so-called weighted SNP correlation network analysis, or WSCNA [51]), which aims to identify clusters of SNPs with similar effects on a given phenotype after controlling for LD. In contrast, we used the WGCNA framework for clustering PRSs across a large number of phenotypes, rather than clustering SNP effects across the genome for a single phenotype.

Extensive details on the WGCNA/WSCNA (v1.71) pipeline have been published previously [48,51]. In our analysis, for each PRS matrix, pairwise Pearson correlations were calculated among all PRSs. For the construction of a weighted network, these PRS-PRS correlations were raised to a soft-thresholding power (*β*), determined empirically using criteria that encourage approximate scale-free topology of the resulting network (via the WGCNA package function *pickSoftThreshold*). We chose to calculate an *unsigned* network, meaning that adjacency was based off of the absolute value of correlation, rather than discarding negative correlations. This was important, as we calculated PRS across thousands of phenotypes, among which both negative and positive correlations would be of interest.

To cluster this network, a topological overlap matrix (TOM) was calculated and used to define dissimilarity for hierarchical clustering. The resulting dendrogram was then clustered using dynamic tree cutting [52] which applies a variable-height tree cut algorithm to maximize the capture of meaningful branch structures (i.e. PRS modules). WGCNA was run using the following additional parameters for detection of PRS modules: module detection cut-height of 0.999, minimum module size of 10, and high branch split sensitivity (deepSplit=4).

Our soft-thresholding power selection varied across PRS matrices (refer to **Figure S3**), depending mostly on the SNP inclusion α-value threshold, and therefore the number of PRSs included in a given matrix. For each matrix, the lowest value of *β* was chosen at which the scale-free topology model fit (R^2^) reached above 0.8 (resulting *β* values ranged from 1 to 6) to satisfy a scale-free topological network. We also chose a low minKMEtoStay parameter (0.01), which governs the behaviour of the dendrogram clustering algorithm and results in the inclusion of nearly all input PRS into some module (rather than excluding the many lowly correlated PRSs from membership in any module). We did this to allow the ePRS pipeline to capitalize on even small genetic influences of weakly-correlated PRSs within clusters, rather than eliminating these subtle contributions entirely. Following the identification of PRS modules, PCA is applied to each module separately, with the first principal component representing that module’s ePRS. These ePRSs were used for downstream modelling of LOAD phenotypes.

### Association analysis of ePRSs with LOAD phenotypes

For each iteration of our PRS analysis (i.e. for each matrix calculated using a different SNP inclusion α-value threshold and including/excluding *APOE* and/or *MHC*), we used multivariate regression models to identify the association of derived ePRSs with the following LOAD-related variables: longitudinal global cognitive decline slope, consensus clinical diagnosis of LOAD at time of death, and brain-wide post-mortem levels of β-amyloid (Aβ) and PHFtau neuropathology. Among the four outcomes, LOAD diagnosis is a binary outcome and was analyzed using logistic regression, with the remaining three continuous traits analyzed using linear models.

For LOAD diagnosis and cognitive decline, regression covariates included biological sex, age at death (age at baseline for decline), and educational attainment. For neuropathological outcomes, educational attainment was replaced with postmortem interval (PMI). False discovery rate (FDR) correction (i.e. q-value [53]) was used to mitigate multiple testing concerns among all ePRS derived from a given matrix; this matrix-specific FDR correction can be thought as a stratified FDR control [54]. For direct comparison with the benchmark PRS from the latest and most powerful GWAS [38] of LOAD, ePRSs demonstrating significant association with at least one of the LOAD phenotypes (q-value<0.05) were carried forward for further analysis.

### Evaluating performance of ePRSs against LOAD PRS benchmark

First, we optimized a single-trait PRS for LOAD (PRS_LOAD_) in the ROS/MAP sample. This was achieved using the same PRSice-based method described previously, calculating PRS_LOAD_ at the 15 different SNP inclusion α-value thresholds, and selecting the top associated (q-value ≤ 0.05) score for each AD-related phenotype. We note that this model selection approach is expected to result in a degree of overfitting for our benchmark PRS_LOAD_, rendering subsequent estimates of ePRS performance gains naturally conservative.

To compare the performance of selected LOAD phenotype-associated ePRSs with our benchmark PRS_LOAD_, we first built two sets of baseline regression models (linear for continuous and logistic for binary outcomes) to establish the performance of covariates alone (Model 1) and covariates + PRS_LOAD_ (Model 2) on our clinical and neuropathological outcomes (*Y*):

**Model 1**. Covariates only:

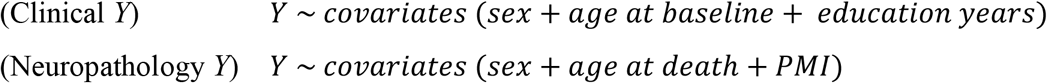

**Model 2**. Covariates and PRS_LOAD_:

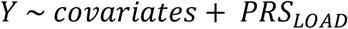

Then, for each ePRS significantly associated with at least one LOAD-related outcome in our preceding analyses, we built a third regression model (Model 3) including covariates, PRS_LOAD_, and the selected ePRS:

**Model 3**. Covariates, PRS_LOAD_, and ePRS:

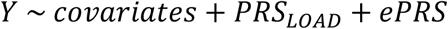

The likelihood ratio test was then used, comparing model 2 with model 3, to assess whether the ePRS significantly improved model fit over PRS_LOAD_ for a given outcome. The difference in model performance was reported as a ΔAUC for logistic and ΔR^2^ for linear models, again comparing models 2 and 3. As each significant ePRS was selected based on association with at least one LOAD-related outcome, we mitigated concerns of overfitting by performing 0.632+ bootstrapping [55] to derive model performance estimates.

## Results

### Principal components analysis of whole-person PRSs in elderly

**Table 1** summarizes demographic information for the individuals included in our study. We first sought to explore the covariance structure of PRSs calculated in our sample for over 2,000 diverse phenotypes when no genomic regions were excluded from our calculations (Π_ALL,α_). **Figure 1** illustrates the overall variation explained by each PC for Π_ALL,α_ (refer to **Figures S4-S6** for the other PRS matrices), and the number of PCs explaining 80% of the total variation of the corresponding matrix. When considering PRSs calculated using a genome-wide significant α-value SNP inclusion threshold of 5 × 10^−8^ (349 PRSs), the first principal component (PC) explained 7.38% of the total variance in all PRSs. In evaluating variable contributions to this PC, we found that the strongest contributing PRSs were those for autoimmune traits, such as chronic hepatitis (total contribution: 3.16%), diffuse diseases of connective tissue (total contribution: 2.98%), Sicca syndrome (total contribution: 2.68%) (**Figure S7**). The LOAD PRS (*PRS*_*LOAD*_) was loaded onto PC28 (total contribution of 2.1% to this PC) at this threshold along with arthrosis-related traits, family history of AD, delirium, dementia, and education, shown in **Figure S7**.

**Figure 1.**
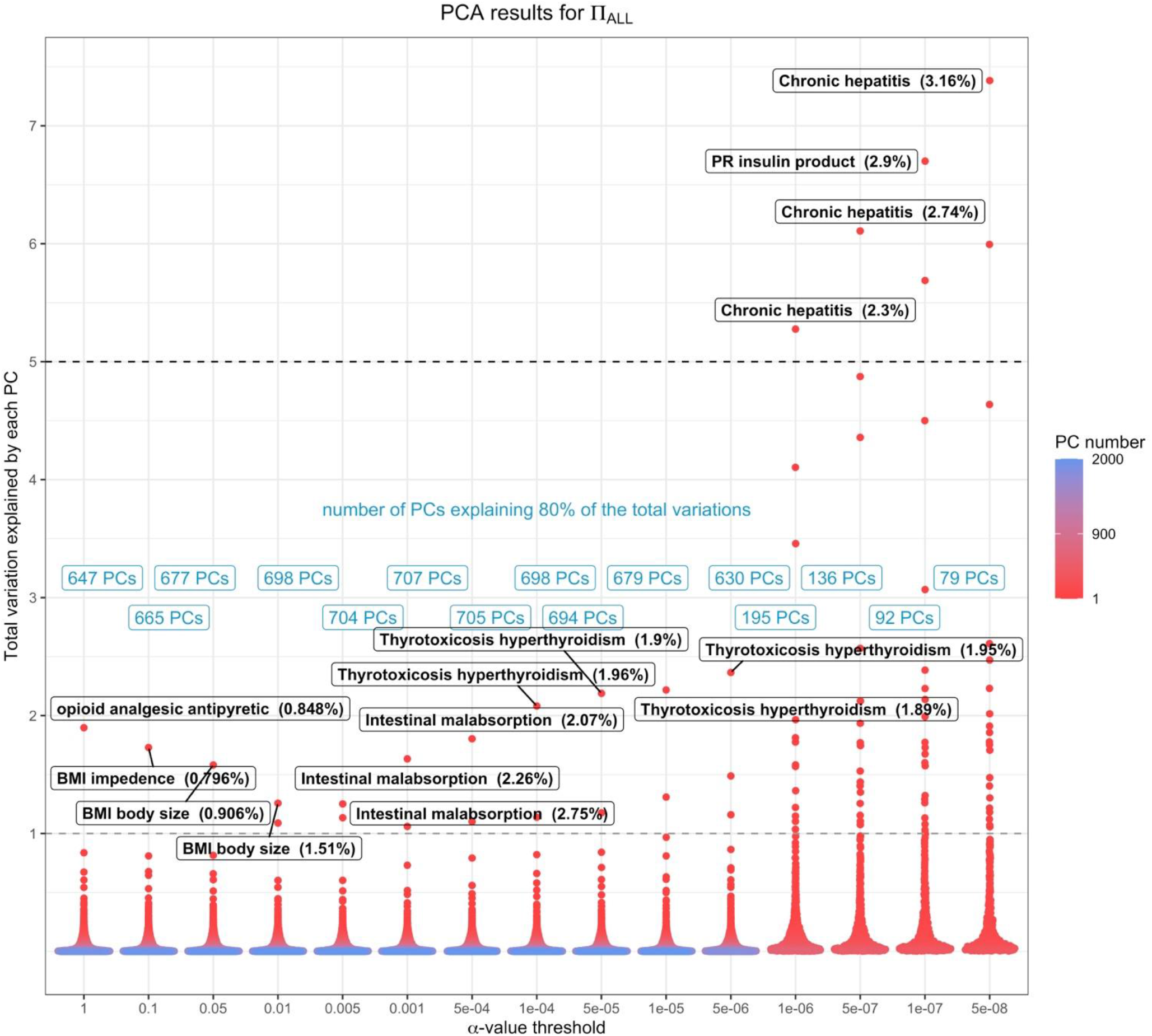
Principal component analysis of whole-person PRS matrices. Beeswarm plot summarizing the results from PCA on *Π*_*ALL*_. Each point represents a PC derived from each PRS matrix, and the colour of each point represents the PC number. The majority of PCs explain less than 1% of the total variation of their corresponding matrices (y-axis). Additionally, a large number of PCs are required to explain 80% of the total variation of the original matrix. However, at lower *α*-value thresholds (x-axis), we notice more PCs explain more than 5% of the total variation of their original matrix, largely due to the smaller number of PRSs at lower *α*-value thresholds. The number one contributing PRS of each PC, with their contributing percentage to the component, is labelled for the first PC of each matrix. **Figures S4-S6** show result summaries for the PCA results on other PRS matrices (*Π*_*MHC*−_, *Π*_*APOE*_ , *Π*_*MHC*/*APOE*−_).

**Table 1.** Demographic summary of Religious Orders Study (ROS) and Memory Aging Project (MAP) participants included in analysis

| Total (n=2,044) | ‡Autopsied (n=1,443) |  |  |  | Not deceased (n=601) |  |  |  |
| --- | --- | --- | --- | --- | --- | --- | --- | --- |
|  | CN (n=473) | MCI (n=345) | LOAD (n=599) | Other (n=26) | CN (n=442) | MCI (n=111) | LOAD (n=47) | Other (n=1) |
| Sex (M/F) | 167/306 | 128/217 | 184/415 | 9/17 | 98/344 | 27/84 | 7/40 | 0/1 |
| Age at study entry (mean (SD)) | 78.77 (6.96) | 81.16 (6.36) | 81.69 (6.74) | 78.48 (7.82) | 73.73 (6.92) | 76.69 (7.33) | 78.17 (8.48) | 87.24 |
| Age at death (mean (SD)) | 87.26 (6.84) | 89.23 (6.31) | 91.22 (5.89) | 88.01 (6.65) | -- | -- | -- | -- |
| §Postmortem interval (mean (SD)) | 9.46 (8.93) | 9.24 (7.56) | 8.52 (6.55) | 10.5 (10.2) | -- | -- | -- | -- |
| Education, years (mean (SD)) | 16.39 (3.70) | 16.12 (3.45) | 16.19 (3.63) | 17.04 (3.90) | 16.70 (3.39) | 15.68 (3.32) | 16.23 (3.10) | 16.00 |
| †Mean LOAD PRS (mean (SD)) | -0.15 (0.79) | -0.03 (0.78) | 0.12 (0.80) | 0.00 (0.86) | -0.03 (0.80) | 0.16 (0.88) | 0.13 (0.80) | 0.13 |

‡191 deceased participants with genotype information did not have quantitative neuropathological data available at time of analysis. §Postmortem interval statistics were calculated on 1,252 people with complete data. †LOAD PRS values shown are averaged over all 15 α-value thresholds within each subgroup (calculated on the whole genome, without removing any regions *a priori*). CN = no cognitive
impairment; LOAD = late-onset Alzheimer's disease; MCI = mild cognitive impairment; Other = cause of dementia not likely Alzheimer's disease (e.g. vascular dementia); SD = standard deviation.

Across all the 15 α-value thresholds, this pattern was fairly consistent, with autoimmune conditions representing the majority of variability in the top PCs. In addition, there was a consistency in the top contributing PRSs to PC1 across all 15 α-value thresholds. At the highest α-value threshold (1.0), PC1 (explaining 1.90% of the variance) was dominated by opioid and generic over-the-counter pain relief medications. Further, PC1 of matrices calculated with somewhat lower α-value threshold values of 0.1, 0.05, and 0.01 (explaining 1.73%, 1.58% and 1.26% of the variance respectively) loaded most prominently onto body mass index (BMI) and weight, along with hypertension and vascular problems. At even lower thresholds of 0.005, 0.001, and 5 × 10^−4^ (explaining 1.25%, 1.63% and 1.80% of the variance respectively) PC1 was primarily represented by PRSs for intestinal malabsorption and hypertension. The PRS for thyrotoxicosis hyperthyroidism was the strongest contributor to PC1 at thresholds of 1 × 10^−4^, 5 × 10^−5^, 1 × 10^−5^ and 5 × 10^−6^ (explaining 2.08%, 2.19%, 2.22% and 2.37% of the variance respectively). PRS of Chronic hepatitis was heavily loaded on to PC1 at α-value threshold of 1 × 10^−6^, 5 × 10^−7^, 1 × 10^−5^ (explaining 5.28% and 6.11% of variance respectively), and lastly usage of insulin loaded onto PC1 at α-value threshold 1 × 10^−7^ explaining 6.70% of the total variance. Detailed breakdowns of PC1 and PC2 at α-value threshold of 0.001 are shown in **Figure S8**.

When the MHC region was removed prior to PRS calculations (Π_MHC−,α_), we found that the top three contributors to PC1 of PRS matrices at lower α-value thresholds (i.e. 5 × 10^−8^, 1 × 10^−7^, 5 × 10^−7^, 1 × 10^−6^, 5 × 10^−6^ and 1 × 10^−5^ explaining 5.17%, 5.03%, 4.78%, 3.81%, 1.21% and 1.02% of the total variance respectively) were mainly influenced by PRSs such as presence of cardiac and vascular implants and grafts, coronary artery bypass grafts, heart attacks, and other heart related problems. PRSs of mean arterial pressure, systolic blood pressure, high blood pressure and hypertension were loaded as top contributors to PC1 at α-value threshold of 5 × 10^−5^, 1 × 10^−4^, 5 × 10^−4^, 0.001 and 0.005 (explaining 0.99%, 0.98%, 1.07%, 1.07% and 1.15% of the total variance respectively). At higher α-value thresholds (α-value of 0.01, 0.05, 0.1), the top PRS contributors to PC1 (explaining 1.25%, 1.57% and 1.71% of the variance respectively) were BMI along with hypertension and other vascular problems. Lastly, PC1 at α-value threshold of 1 was similar to the PC1 calculated with the whole genome, and it consisted of PRSs of opioid use, and other over the counter pain killers. More detailed description of PC1 and PC2 at α-value threshold of 0.001 are shown in **Figure S9**.

A few of the PCs were heavily influenced by LOAD, dementia, and family history of AD with other known AD-related traits such as high cholesterol, usage of statin, apolipoprotein biomarker, directly assessed levels of LDL cholesterol (LDL) and delirium. Though these PCs were mainly present in Π_ALL,α_ and Π_MHC−,α_. Additionally, the contribution of PRS of LOAD was low compared to the top contributing PRSs to these PCs. Interestingly, PRS of LOAD was not present in AD-related PRSs PC when the MHC region was not included. **Figures S7, S10 and S11** illustrate a few selected LOAD related PCs from Π_ALL,α_ and Π_MHC−,α_. In the absence of the *APOE* region (Π_APOE−,α_), there was little notable change in the variability explained by PCs or in the structure of those PCs (when compared to Π_ALL,α_). This was true across all α-value thresholds, with a few exceptions. PC1 at α-value thresholds of 5 × 10^−8^ (350 PRSs, explaining 7.32% of the total variations) influenced autoimmune traits like chronic hepatitis (contribution of 3.16%), Thyrotoxicosis hyperthyroidism (contribution of 2.99%), and Sicca syndrome (contribution of 2.78%). In the absence of both influential genomic loci (Π_MHC/APOE−,α_) PC1 at α-value thresholds of 5 × 10^−8^ (315 PRSs, explaining 5.12% of the total variance) was mostly influenced by coronary artery bypass grafts (contribution of 4.96%). Lastly, the role of *PRS*_*LOAD*_ in contributing to the PCs from both Π_APOE−,α_ and Π_MHC/APOE−,α_ dropped notably.

### Clustering of PRS matrices using WGCNA

Following descriptive PCA and the assessment of phenome-wide variance in PRSs at different α-value thresholds, we sought to group PRSs into discrete clusters based on their network-based similarity using WGCNA. Unlike PCA, WGCNA provides discrete, mutually exclusive groupings of PRS subnetworks for further analysis. Beginning with the entire genome and an α-value threshold of 0.05 (Π_ALL,0.05_), which includes a large number of SNPs per phenotype, the WGCNA approach yielded 33 PRS clusters (or modules). As expected, biologically and clinically related traits clustered together. The number of modules identified, differed as a function of α-value threshold (12 modules from 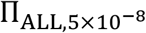, 15 modules from 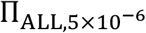, 15 modules from 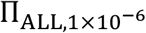, 35 modules from Π_ALL,1_, the rest are shown in **Table S3**), though module identities remained largely cohesive. Hub analysis was used to reveal the PRSs that were most interconnected within each module; hub PRSs at different thresholds included those for weight in the red module (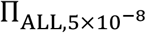, consisting of 22 PRSs), rheumatoid arthritis in the blue module (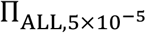, consisting of 209 PRSs), and malignant melanoma in the black module (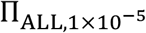, consisting of 25 PRSs), and cancer of bronchus lung in the brown module (Π_ALL,0.01_, consisting of 103 PRSs). Note that module color labels are arbitrarily assigned by WGCNA and not linked to particular sets or sizes of PRS (i.e. they are not preserved between runs of WGCNA). Some modules, such as the weight module, were largely preserved across nearly all α-value (except for 1 × 10^−6^ and 5 × 10^−7^) thresholds (**Figures S12-S15**); this module was clustered with PRSs related to body mass, bone density, and coronary and heart diseases. Conversely, chronic ischemic heart disease in the yellow module (*Π*_*ALL*,5e−05_, consisting of 21 PRSs) clustered along with other heart or coronary complications such as heart attack (myocardial infarction) and ischemic heart disease. Notably, this module was conserved across α-value thresholds of 0.01, 0.005, 5 × 10^−5^ and 5 × 10^−7^, and was correlated with modules that included PRS_LOAD_ and PRSs for family history of AD and all-cause dementia. Of note, PRS_LOAD_ consistently clustered within the rheumatoid arthritis module in Π_*ALL*,0.001_, Π_*ALL*,0.0005_, Π_*ALL*,0.0001_, *Π*_A*POE*−,0.001_, *Π*_A*POE*−,0.0005_, *Π*_*APOE*−,0.0001_, 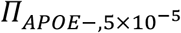. **Figure S16** outlines the correlational structure of two selected modules (weight and high cholesterol).

Focusing specifically on the PRS_LOAD_, we found it clustered within the pink module from *Π*_*ALL*,5e−08_ (consisting of 19 PRSs), red module from *Π*_*ALL*,1e−07_ (consisting of 23 PRSs) and Π_A*LL*,5e−05_ (consisting of 22 PRSs), brown module from *Π*_*ALL*,1e−06_ (consisting of 68 PRSs) and *Π*_*ALL*,5e−06_ (consisting of 85 PRSs) and *Π*_*ALL*,1e−05_ (consisting of 77 PRSs) along with PRSs for all-cause dementia, family history of AD or AD-dementia, delirium, Apolipoprotein levels, cholesterol levels, presence of cardiac and vascular implants and grafts – reinforcing existing evidence for genetic correlation between these traits in an unbiased manner. History of taking prescriptions to treat vascular and abnormal cholesterol levels, such as rosuvastatin, ezerimibe, and HMG-CoA reductase inhibitor statins, also clustered with PRS_LOAD_ and other dementias at lower α-value thresholds. Interestingly, the PRS_LOAD_ did not consistently cluster with the UKB-derived family history of AD dementia (ICD code: G30.9), family history of AD (ICD code: G30.9) or dementia (ICD code: F03), PRSs – demonstrating the divergence of PRS derived from different cohorts. However, this divergence was only present at higher α-value thresholds (*Π*_*ALL*,1_, *Π*_*ALL*,0.01_, *Π*_*ALL*,0.05_, *Π*_*ALL*,0.01_) reinforcing the importance of SNP inclusion when using clumping and thresholding [40].

The removal of MHC and/or *APOE* loci resulted in several changes in module number and membership across α-value thresholds. Most notably, when the *APOE* region was excluded (*Π*_*APOE*−,α_), the PRS_LOAD_ clustered either in the grey module (this module contains a set of PRSs which have not been clustered in any module), or the turquoise module (this module contains the highest number of PRSs), along with other dementia-related phenotypes (and many others). This was true across all α-value thresholds except for the blue module (*Π*_*APOE*−,0.01_ represented by weight, *Π*_*APOE*−,0.005_ represented by BMI,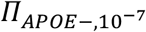 represented by family history of diabetes, 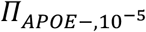 represented by prednisolone, 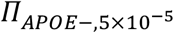 represented by Rheumatoid Arthritis (RA)), yellow module (*Π*_*APOE*−,0.001_ represented by RA), and the brown module (*Π*_*APOE*−,0.0005_ represented by RA) where the PRS_LOAD_ was clustered with other PRSs of which none were PRS of dementia or family history of AD. When the MHC region was also excluded, there were some modules where PRS_LOAD_ was clustered with PRS of dementia or family history of AD or delirium. However, this membership was not consistent and varied for different α-value thresholds. Furthermore, PRS_LOAD_ appeared in the blue module (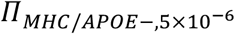 represented by presence of cardiac and vascular implants and grafts, *Π*_*MHC*/*APOE*−,0.01_ represented by weight).

### Associations of ePRS with clinical and neuropathological traits

Following the network-based identification of correlated PRSs across the phenome, we evaluated the associations of derived ePRSs with late-life cognitive and neuropathological phenotypes. **Figure 2** summarizes these results across all *α*-value thresholds (and all PRS matrices), highlighting several modules strongly associated with our LOAD-related phenotypes at a false discovery rate (FDR) adjusted q-value ≤ 0.05. Notably, all significant ePRS associations were observed for the Π_ALL,α_ and Π_MHC−,α_ matrices, whereas none of the ePRSs calculated from Π_APOE−,α_ and Π_MHC−APOE−,α_ passed FDR correction. Full sets of ePRS association results are summarized in **Figures S18-S20**.

**Figure 2.**
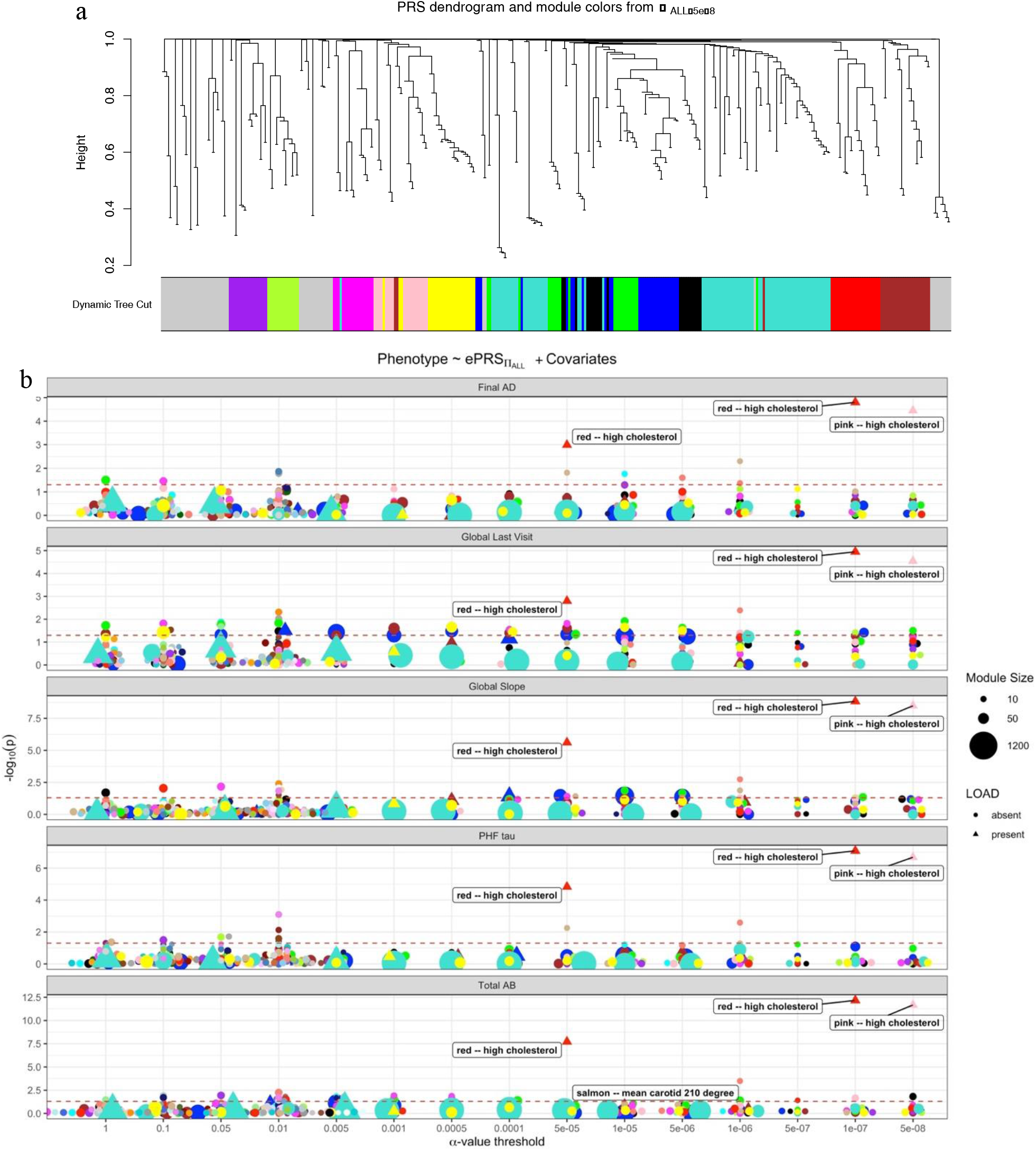
Eigen-PRS associations with LOAD-related phenotypes. **(A)** selected dendrogram (based on the WGCNA topological overlap matrix of *Π*_*All*,5*e*−06_) produced by the WGCNA pipeline. **B)** Dotplots summarizingthe associations of ePRSs derived from *Π*_*All*_ with each LOAD phenotype. Hub PRSs for significantly associated ePRSs (FDR q-value ≤ 0.05) are labelled. The red dotted line represents uncorrected *p*=0.05. odule represented by high cholesterol (pink) at α-value threshold of 5 × 10^−7^ had the highest association with all five phenotypes. The modules, represented by high cholesterol levels (pink, purple) showed the strongest associations with all five aging phenotypes at α-value threshold of 1 × 10^−6^, 1 × 10^−7^, and 5 × 10^−8^. A triangular dot shape indicates that the LOAD PRS calculated from Bellenguez et al. (2022) was a member of the PRS cluster from which a given ePRS was calculated. Refer to **Figure S18, S19, and S20** for summary results from PRS matrices from sensitivity analyses (*Π*_*MHC*−_, *Π*_*APOE*−,_, *Π*_*MHC*−*APOE*−_). 20

Among the top results were the pink ePRS from *Π*_*All*,5e−08_, the red module from Π_A*ll*,1e−07_, and the red module from *Π*_*All*,5e−05_, *Π*_*All*,1e−06_, which were each largely composed of PRSs for cholesterol levels, and strongly associated with all five LOAD phenotypes. EPRSs with hub PRSs of maximum carotid intima-medial thickness (IMT; based on ultrasound) were strongly associated with β-amyloid levels. With the removal of the MHC region, the top associated ePRSs with all five phenotypes were consistent in terms of overall PRSs in these modules. Though, more ePRSs represented by high cholesterol indicated association with all five phenotypes. Overall, ePRSs represented by hubs of high cholesterol levels, LDL, and maximum carotid IMT were found to be highly associated with the LOAD-related traits and neuropathologies.

### Assessment of improvements in predictive performance of ePRS over established single-trait LOAD PRS

Having identified several ePRSs associated with LOAD-related traits and neuropathologies, we sought to test whether these ePRSs could significantly augment single-PRS models of only PRS_LOAD_ and basic demographics. In some cases, the ePRSs were able to improve the variance explained by the base models by up to 1.7%. Modelling results for selected top ePRSs are shown in **Table 2**. The brown hub of *Π*_*MHC*−,1e−05_, represented by the hub PRS for cholesterol levels, was able to improve predictive models over the single-trait PRS_LOAD_ for both β-amyloid counts (*P*_*likelihood*−*ratio test*_ = 5.29 × 10^−5^) and global random slope of cognitive decline (*P*_*likelihood*−*ratio test*_ = 3.16 × 10^−3^). Similarly, the red module, represented by high cholesterol levels, from *Π*_*All*,1e−07_, was able to improve the predictability power of the PRS_LOAD_ in association with β-amyloid count (*P*_*likelihood*−*ratio test*_ = 9.25 × 10^−6^). **Figure 3** (**expanded in S21**) summarizes the overall performance comparison between base models and the ePRS-inclusive full models (defined as model 2 and model 3). Only ePRSs with FDR-adjusted q-value ≤ 0.05 for the likelihood ratio test between base and full model are shown.

**Figure 3.**
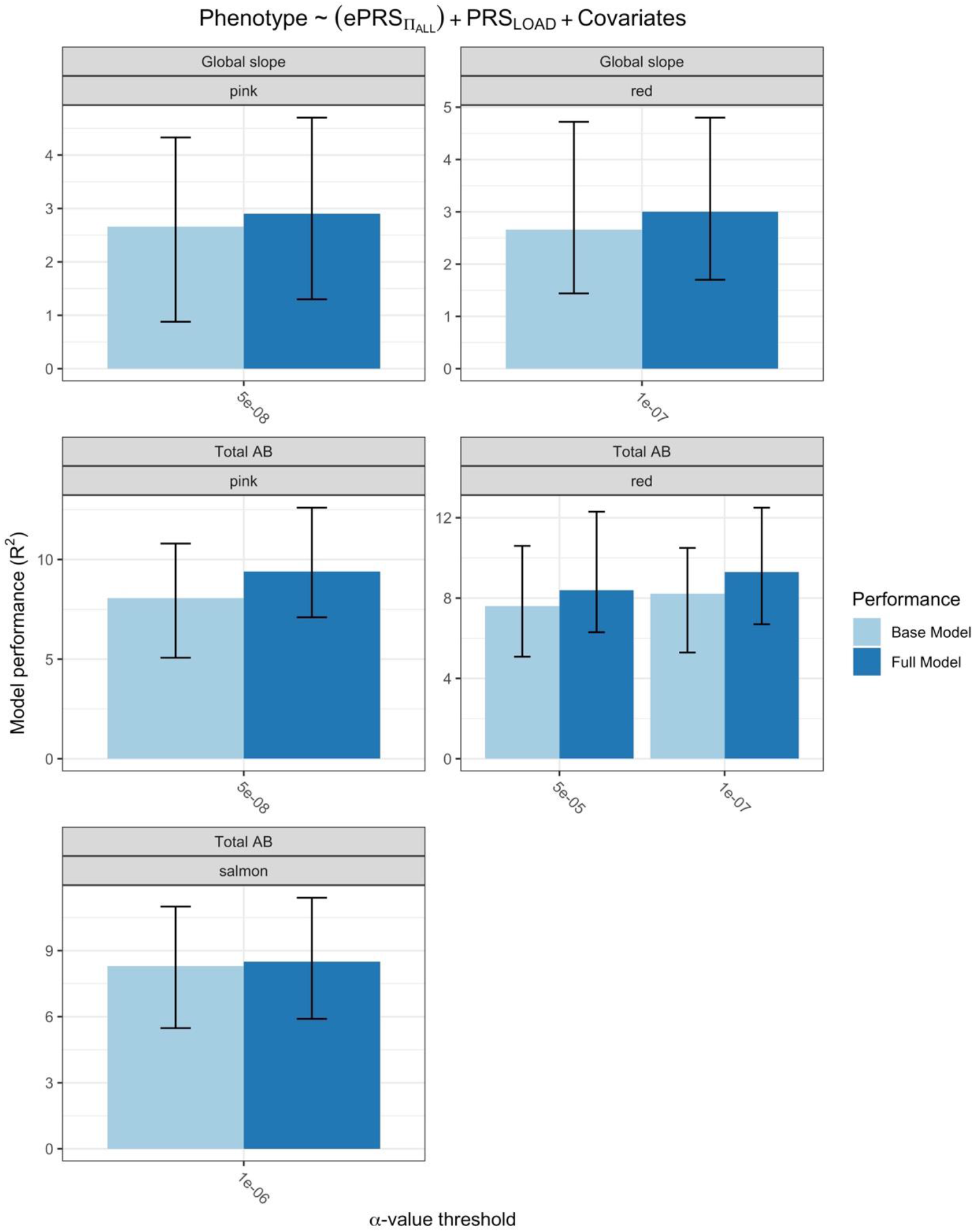
Summary of model improvements for ePRS-inclusive models over single-trait LOAD PRS models of LOAD-related phenotypes. Barplots showing the results of the comparison of full models including ePRSs and single-trait PRS models of PRS_LOAD_, calculated from Π_*ALL*,α_. For continuous response variables (Aβ burden, PHF tau, cognitive global random slope, cognitive global random slope at last visit) we report the .632 bootstrapped variation explained (*R*^2^) by the covariates for the base (model 2) and full model (model 3). Results for ePRS derived from matrices excluding the MHC region are shown in **Figure S21**. Error bards shown are estimates of 95% confidence intervals for variance explained with bootstrapping. Significance of differences in model performance were determined using likelihood ratio tests between the base and full models prior to bootstrapping, followed by FDR correction (all shown results had corrected *q*-value < 0.01).

**Table 2.** Evaluation of significant ePRSs models against standard single-trait PRS_LOAD_ models of Alzheimer’s neuropathology and cognitive decline.

| PRS Matrix | Outcome | ePRS<br>(module) | $\beta_{ePRS}$ | Model<sup>##</sup><br>Performance<br>(95% CI) | ΔModel<sup>###</sup><br>performance<br>$P_{LRT}$ | n |
| --- | --- | --- | --- | --- | --- | --- |
| $\Pi_{All, 5 \times 10^{-5}}$ | Total Aβ | Red<br>(High cholesterol) | 5.105 | 0.084<br>(0.063,0.123) | 0.008<br>$4.69 \times 10^{-4}$ | 1238 |
| $\Pi_{All, 1 \times 10^{-6}}$ | Total Aβ | Salmon<br>(Mean Carotid) | 3.634 | 0.085<br>(0.059,0.114) | 0.002<br>$8.63 \times 10^{-3}$ | 1238 |
| $\Pi_{All, 1 \times 10^{-7}}$ | Global slope | Red<br>(High Cholesterol) | -0.334 | 0.030<br>(0.017,0.048) | 0.003<br>$1.11 \times 10^{-3}$ | 1915 |
| | Total Aβ | | 6.770 | 0.093<br>(0.067,0.125) | 0.011<br>$9.25 \times 10^{-6}$ | 1238 |
| $\Pi_{All, 5 \times 10^{-8}}$ | Global slope | Pink<br>(High Cholesterol) | -0.325 | 0.029<br>(0.013,0.047) | 0.002<br>$1.46 \times 10^{-3}$ | 1915 |
| | Total Aβ | | 6.545 | 0.094<br>(0.071,0.126) | 0.013<br>$1.57 \times 10^{-5}$ | 1238 |
| $\Pi_{MHC-, 0.0001}$ | Total Aβ | Green<br>(High Cholesterol) | 0.179 | 0.091<br>(0.073,0.129) | 0.017<br>$9.02 \times 10^{-7}$ | 1238 |
| $\Pi_{MHC-, 5 \times 10^{-5}}$ | Global slope | Yellow<br>(High Cholesterol) | -0.017 | 0.093<br>(0.071,0.121) | 0.013<br>$1.10 \times 10^{-5}$ | 1915 |
| | Total Aβ | | 0.181 | 0.091<br>(0.059,0.121) | 0.010<br>$2.22 \times 10^{-5}$ | 1238 |
| $\Pi_{MHC-, 1 \times 10^{-5}}$ | Global slope | Brown<br>(High Cholesterol) | -0.015 | 0.085<br>(0.061,0.119) | 0.005<br>$3.16 \times 10^{-3}$ | 1915 |
| | Total Aβ | | 0.163 | 0.085<br>(0.064,0.122) | 0.012<br>$5.29 \times 10^{-5}$ | 1238 |
| $\Pi_{MHC-, 1 \times 10^{-6}}$ | Global slope | Brown<br>(High Cholesterol) | -0.013 | 0.030<br>(0.015,0.048) | 0.004<br>$9.71 \times 10^{-4}$ | 1915 |
| | Total Aβ | | 0.180 | 0.079<br>(0.054,0.118) | 0.008<br>$3.39 \times 10^{-4}$ | 1238 |
| $\Pi_{MHC-, 5 \times 10^{-7}}$ | Global slope | Red<br>(High Cholesterol) | -0.013 | 0.036<br>(0.024,0.053) | 0.004<br>$2.02 \times 10^{-3}$ | 1915 |
| | Total Aβ | | 0.179 | 0.028<br>(0.013,0.048) | 0.002<br>$1.88 \times 10^{-3}$ | 1238 |
| $\Pi_{MHC-, 1 \times 10^{-7}}$ | Total Aβ | Black<br>(LDL) | 0.205 | 0.084<br>(0.058,0.131) | 0.003<br>$4.16 \times 10^{-3}$ | 1238 |
| $\Pi_{MHC-, 5 \times 10^{-8}}$ | Total Aβ | Green<br>(LDL) | 0.209 | 0.037<br>(0.023,0.052) | 0.001<br>$1.37 \times 10^{-2}$ | 1238 |

Model performance is provided as variance explained ( $R^2$ ) after .632 bootstrapping, including 25% confidence interval. For $\Delta$ model performance, the $\Delta R^2$ value is shown as well as the $p$ -value of the likelihood ratio test between the full and base models. \* $p$ = <0.05, \*\* $p$ = <0.01, \*\*\* $p$ = 0.001. All $p$ -values are reported at an uncorrected level. Global slope = slope of change in longitudinal global cognitive performance; n = sample size.

## Discussion

We generated over 120,000 PRSs from 2,218 phenotypes in a sample of over 2,000 elderly individuals harnessing the latest GWAS summary statistics from the Pan-UK Biobank Consortium. We used a network-based approach to derive clusters of PRSs and identified meaningful patterns of biologically and clinically related phenotypes with relevance to cognitive decline and core LOAD-related neuropathologies. By including summary statistics for thousands of diagnoses, traits, biomarkers, behaviours, and health-related measures, rather than selecting major risk factors *a priori*, we build a richer understating of the relationship between the genetic structure of our elderly population and risk for LOAD.

In PCA-based exploration of the full set of PRSs at each alpha threshold, loadings on the top PCs uncovered both previously observed genetic correlations of certain phenotypes, as well as some new ones; at lower α-value thresholds, corresponding to genome-wide and near genome-wide significant SNPs only, the top PCs were largely dominated by PRSs for cholesterol levels and cardiovascular related health conditions, treatments, and procedures. At higher α-value thresholds, we found PRSs for pain medications, including opioids and general over-the-counter painkillers, such as paracetamol, dominating the loadings of top PCs. We hypothesize that this is due to the wide use of these medications for broad spectra of clinical and subclinical afflictions, and therefore genetic associations with their use would tag variation common across hundreds or thousands of included phenotypes (from minor injury risk, to cancers, to stress-related events and conditions). Unsurprisingly, top PCs explained smaller proportions of variance across PRS matrices that included more PRSs (i.e. those calculated using less stringent, higher alpha thresholds). However, the proportion of PCs (out of number of included PRSs) required to account for 80% of variance in all PRSs was largely consistent across larger *α*-value thresholds (5 × 10^−6^ to 1). Compared to similar work [21], the LOAD PRS consistently clustered PRS for cholesterol levels and statin prescription, with both PRSs contributing highly to their corresponding PC. However, we also noticed the influence of apolipoprotein biomarker and LDL in these PCs as well. While PCA provided useful insight into the primary axes of variability in our PRS matrices, the utility of the resulting PCs for genetic prediction are not clear; for modelling there are no reliable guidelines for which PCs to include at each *α*-value threshold, and the variation explained by the top PCs was less than 1% for the majority of the PCs.

Thus, we applied a clustering method which is traditionally used on gene co-expression data, WGCNA. In most cases, ePRSs derived from the WGCNA pipeline that were significantly associated with LOAD phenotypes were composed of a combination of PRSs for dementia-related phenotypes (e.g. family history of LOAD, LOAD diagnosis, and all-cause dementia), cholesterol levels, prescriptions for statins and other cardiovascular health medications, diseases, and physiological measurements of carotid arteries. This reinforces the known links between cardiovascular health and risk for LOAD [56], but also points toward the specific aspects of cardiovascular health that may be more genetically important for determining this risk than others. In sensitivity analyses removing the *APOE* and/or MHC regions from our scores, we noted major impacts on the associations of derived ePRS on LOAD phenotypes. In fact, we found no associations (q-value < 0.05) between ePRSs produced from *Π*_*APOE*−_ and *Π*_*MHC*/*APOE*−_ and any LOAD phenotypes. Even ePRSs that included PRS_LOAD_ were not associated with LOAD-related phenotypes, though interestingly PRS_LOAD_ did cluster with PRSs for rheumatoid arthritis, which has been shown to be genetically related to microglial activation that is implicated in LOAD pathogenesis [32].

This study has several limitations. First, our conclusions related to the relationship between heritable risk for our phenotypes of interest are based on summary statistics derived from the UK Biobank (indeed, those of any PRS-based study). The UK Biobank has known healthy volunteer bias [57], and our target ROS/MAP sample also has a known late-life resilience bias as a result of survival to study entry without dementia and lifestyle factors associated with part of its underlying religious population [25]. Therefore, the transfer of genetic risk profiles from one population to the other may be subject to challenges with cross-sample portability – most apparent when considering ancestral differences but extend to age, sex, and measurement differences between groups - that may hide true genetic effects [58]. This issue of portability is also made clear by the observation that our PRS_LOAD_ did not always cluster with UKB summary stats-derived LOAD and dementia PRSs; in part likely due to differences in phenotypic definition between studies. Second, by clustering matrices across traits and within SNP inclusion thresholds, we have assumed that meaningful correlations between PRSs may not occur between different traits at different thresholds. Others have attempted to resolve this, for example by calculating the first principal component of a set of PRSs derived from the same set of summary statistics across multiple thresholds [42]. However, we sought specifically to find relationships between the PRSs rather than finding the most optimized α-value threshold of each PRS, since it would require availability of the same phenotypes in the target population. Further, while we used a 5% threshold on estimated heritability in the UK Biobank to ensure inclusion of only traits with a nonzero genetic basis, there is no guarantee that PRS models based on these GWAS will have a significant predictive power. Therefore, we acknowledge that our observed clusters of PRSs may have been biased by patterns of unquantified within-trait predictive performance, which could only be estimated at scale in fully-phenotyped (i.e. all 2000+ traits) out-of-sample data. Methods have been proposed for addressing this limitation in PRS tuning [59] and they will be explored in future work.

Finally, we selected only sex non-specific phenotypes to generate their corresponding PRSs using commonly used methods. Thus, our analysis omits information available for sex-specific traits, such as those related to menopause or testicular cancers, which are known to impact aging in important ways. Additionally, evidence suggests sex-specific effects of LOAD genetic risk factors in the clinical progression of LOAD and the accumulation of LOAD-related neuropathologies [60]; these effects were not explored here and are the focus of ongoing work.

In sum, our study provides a whole-person view of the genetic risk landscape for LOAD in an elderly population. We highlight the impacts of 1) varying SNP inclusion *α*-value thresholds on this landscape using the most popular PRS calculation method, 2) the links between several blood-based and ultrasound imaging-derived cardiovascular traits on LOAD risk, as well as 3) the important contribution of the *APOE* region. Our results suggest that combining genetic risk profiles for vascular and dementia-related phenotypes using latent methods may offer improvements over existing single-trait PRS models of LOAD, a step toward achieving the unrealized potential of PRS for risk stratification in populations and clinics.

## Supporting Information

The Supporting Document contains **Supplementary Methods, Tables S1-S3** (**Table S1** is stored as an external CSV file due to its size), and **Figures S1-S20**.

### Declaration of Interests

The authors declare no competing interests.

## Acknowledgements

DF is supported by the Michael and Sonja Koerner Family New Scientist Award, the Krembil Family Foundation, the Canadian Institutes of Health Research (CIHR), and the Centre for Addiction and Mental Health (CAMH) Discovery Fund. LS is supported by The Natural Sciences and Engineering Research Council of Canada (NSERC), RGPIN-04934 and RGPAS-522594. ROS/MAP is supported by P30AG10161, P30AG72975, R01AG15819, R01AG17917. U01AG46152, U01AG61356.

## Data and Code Availability

All phenotypes and omics data of the Religious Orders Study (ROS) and Memory Aging Project (MAP) Cohorts are available through the RADC Resource Sharing Hub at www.radc.rush.edu. ROS/MAP resources can be requested at https://www.radc.rush.edu. The UK Biobank GWAS summary statistics are available from Neale Lab, http://www.nealelab.is/ukbiobank. All scripts used for this manuscript are open-source and available at https://github.com/aminekhasteh/ePRS.

## Notes

### Competing Interest Statement

The authors have declared no competing interest.

### Summary of Updates

Addition of supplementary information.

