## Supplementary Information for "Association of Whole-Person Eigen-Polygenic Risk Scores with Alzheimer’s Disease"

### LIST OF TABLES

S1 In this table we outline the selected GWAS summary statistics of phenotypes used in this study. The GWAS summary statistics were taken from the Pan-UK biobank consortium, except for GWAS of LOAD which was taken from [1]. Each column is previously defined by the Pan-UK biobank (except for LOAD). Phenocode is the code used for the phenotypes, and trait type is one of the following: continuous, biomarkers, prescriptions, icd10, phecode, categorical. The phenotype name is the name of each phenotype as they appear in this study. ICD10 is the coding of the phenotypes defined by the International Classification of Diseases. Cases and control represent the number of case and control (only European population) in each GWA study. Heritablity is the saige heritablity measure of each GWAS (except for LOAD). $\lambda_{GC}$ is the genomic control of each GWAS (except for LOAD). Lastly the last column provides the link to each summary statistics. . . . 6

S2 Total number of PRSs before (before QC) and after (after QC) applying initial quality control. At each $\alpha$-value threshold (first column), we removed PRSs which were calculated with less than 5 SNPs. We also removed duplicated PRSs, as described in methodology. We did this process across all PRS matrices ($\Pi_{ALL,\alpha}$, $\Pi_{MHC-,\alpha}$, $\Pi_{APOE-,\alpha}$, $\Pi_{MHC/APOE-,\alpha}$). . . . 8

S3 Here you will find a list of the number of modules selected by the WGCNA based on all $\alpha$-value thresholds and genome region inclusion criteria. The highest number of modules was 62, constructed from $\Pi_{APOE-,\alpha \leq 0.01}$. . . . . 16

### LIST OF FIGURES

S1 Distribution of postmortem pathology and rates of antemortem cognitive decline calculated in n=2,044 participants from the Religious Orders Study and Memory and Aging Project (ROS/MAP). **A**. $\beta$-Amyloid protein (n=1,238) identified by molecularly-specific immunohistochemistry and quantified by image analysis. Value is percent area of cortex oocupied by amyloid beta. Mean of $\beta$-Amyloid score in 8 regions [2]. **B**. Neuronal neurofibrillary tangles (n=1,247) are identified by molecularly specific immunohistochemistry (antibodies to abnormally phosphorylated Tau protein, AT8). Cortical density is determined using systematic sampling. Mean of tangle score in 8 regions (4 or more regions are needed to calculate) [2]. **C**. Measurement of global cognitive decline (n=1,915). **D**. Last measurement of global cognitive decline (n=2,040). . . . . 5

S2 Multidimensional scaling (MDS) plot of 1000 Genome against the ROS/MAP dataset 2,067 individuals. The black crosses (+='OWN') in the upper left part represent the first two MDS components of the individuals in the ROS/MAP dataset (The colored symbols represent the 1000 Genome data ('EUR'=European; 'AFR'=African; 'AMR'=Ad Mixed American; 'ASN'=Asian)). The MDS components representing the European samples (the green color) is overlapped with our dataset, showing the homogeneity of our population. Each figure S2(a,b) are showing the MDS plot in different dimensions. . . . . 7

S3 To achieve Scale-Free Topology, we select the smallest $\beta$ value so that the $R^2$ of equation S14 is highest. The $\beta$ values selected in constructing the Scale-Free Topology are labelled in this plot. As $\beta$ increases, we have smaller mean connectivity (refer to equation S13). Additionally, we notice that with higher $\alpha$-value threshold, $\beta = 1$, indicating that by including every gene in our PRS calculations for all phenotypes and the 2,044 ROS/MAP participants, we obtain a scale-free topology. . . . . 10

S4 Principal Component Analysis of phenome-wide PRS. This plot outlines the results from the PC analysis on $\Pi_{MHC-}$. Each point is a component for each matrix, and the colour of each point represents the PC number. The top contributor of each PC, with their contributing percentage to the component is shown for the first PC of each matrix. These PCs are mainly dominated by autoimmune diseases at lower $\alpha$-value thresholds, and general pain medications at higher $\alpha$-value thresholds. . . . . 11

S5 Principal Component Analysis of phenome-wide PRS. This plot outlines the results from the PC analysis on $\Pi_{APOE-}$. Each point is a component for each matrix, and the colour of each point represents the PC number. The top contributor of each PC, with their contributing percentage to the component is shown for the first PC of each matrix. These PCs are mainly dominated by autoimmune diseases at lower $\alpha$-value thresholds, and general pain medications at higher $\alpha$-value thresholds. . . . . 12

S6 Principal Component Analysis of phenome-wide PRS.This plot outlines the results from the PC analysis on $\Pi_{MHC/APOE}$. Each point is a component for each matrix, and the colour of each point represents the PC number. The top contributor of each PC, with their contributing percentage to the component is shown for the first PC of each matrix. These PCs are mainly dominated by autoimmune diseases at lower $\alpha$-value thresholds, and general pain medications at higher $\alpha$-value thresholds. . . . . 13

S7 Top 25 contributors to PC1 and PC28, calculated from $\Pi_{ALL}$, are shown with their contribution. Each dot represents 0.4% of the PRS contribution to this PC. . . . . 17

S8 Top 25 contributors, with their contribution, to PC1 and PC2, calculated from $\Pi_{ALL,0.001}$, are shown here. Each dot represents 0.4% of the PRS contribution to this PC. . . . . 18

S9 Top 25 contributors, with their contribution, to PC1 and PC2, calculated from $\Pi_{MHC-,0.001}$, are shown here. Each dot represents 0.4% of the PRS contribution to this PC. . . . . 19

S10 Top 25 contributors, with their contribution, to PC12 and PC9, calculated from $\Pi_{ALL,1e-05}$ and $\Pi_{ALL,5e-08}$, are shown here. Each dot represents 0.4% of the PRS contribution to this PC. . . . . 20

S11 Top 25 contributors, with their contribution, to PC6 and PC10, calculated from $\Pi_{MHC-,5e-07}$ and $\Pi_{MHC-,1e-05}$, are shown here. Each dot represents 0.4% of the PRS contribution to this PC. . . . . 21

S12 Module preservation of the PRSs at every $\alpha$-value threshold for $\Pi_{ALL}$. On the y-axis, we present the most interconnected PRS in each module (shown as circles or triangles). The modules are outlined by their corresponding module name, and modules that include $PRS_{LOAD}$ are shown as triangles. We only included modules that are preserved at least two times across all 15 $\alpha$-value threshold. The rest of the modules are ordered from most conserved module (Weight) to least. . . . . 22

S13 Module preservation of the PRSs at every $\alpha$-value threshold for $\Pi_{MHC-}$. On the y-axis, we present the most interconnected PRS in each module (shown as circles or triangles). The modules are outlined by their corresponding module name, and modules that include $PRS_{LOAD}$ are shown as triangles. We only included modules that are preserved at least two times across all 15 $\alpha$-value threshold. The rest of the modules are ordered from most conserved module (mean carotid) to least. 23
S14 Module preservation of the PRSs at every $\alpha$-value threshold for $\Pi_{APOE-}$. On the y-axis, we present the most interconnected PRS in each module (shown as circles or triangles). The modules are outlined by their corresponding module name, and modules that include $PRS_{LOAD}$ are shown as triangles. We only included modules that are preserved at least two times across all 15 $\alpha$-value threshold. The modules are ordered from most conserved module (Weight) to least. With the removal of the *APOE* region, we did not identify a module mostly interconnected with family history of AD dementia or any other AD-related PRS. . . . . . . . . . . . . . . . . . . . . . . . . . . . . . . . . . . . . . . . . 24
S15 Module preservation of the PRSs at every $\alpha$-value threshold for $\Pi_{MHC/APOE}$. On the y-axis, we present the most interconnected PRS in each module (shown as circles or triangles). The modules are outlined by their corresponding module name, and modules that include $PRS_{LOAD}$ are shown as triangles. We only included modules that are preserved at least two times across all 15 $\alpha$-value threshold. The modules are ordered from most conserved module (mean carotid) to least. With the removal of the *APOE* region, we did not identify a module mostly interconnected with family history of AD dementia or any other AD-related PRS. . . . . . . . . . . . . . . . . . . . . . . . . . . 25
S16 Outlining the correlational structure of the hub PRSs for weight and high cholesterol across all $\alpha$-value thresholds for all 4 PRS matrices. We notice that in both heatmaps, the hub PRSs are highly correlated with each other, verifying the robustness of these modules. . . . . . . . . . . . . . . . . . . . 26
S17 This plot presents the total variation explained by PC1 in each module (generated by WGCNA). This measurement is highly dependent on the size of the module. The modules in the higher thresholds consist of a larger number of PRSs, explaining lower variance. Conversely, we can see smaller-sized modules at lower $\alpha$-value thresholds explaining higher variance (more than 40% on average). We identified modules, where their corresponding PC1 captured more than 60% of the total variance. . 27
S18 Summarising the association of the aging phenotypes from the ROS/MAP study cohort and the ePRSs from $\Pi_{MHC-}$. The red dotted line represents 5% significance level. We outline the hub PRS of the highly associated module with the ageing phenotypes (q-value $\leq$ 0.05). These modules were selected to test if they can improve the predictability ability of state-of-the-art $PRS_{LOAD}$. . . . . . . . 28
S19 Summarising the association of the aging phenotypes from the ROS/MAP study cohort and the ePRSs from $\Pi_{APOE-}$. The red dotted line represents 5% significance level. We outline the hub PRS of the highly associated module with the ageing phenotypes (smallest p-value, no module passed q-value $\leq$ 0.05 testing). These modules were selected to test if they can improve the predictability ability of state-of-the-art $PRS_{LOAD}$. . . . . . . . . . . . . . . . . . . . . . . . . . . . . . . . . . . . 29
S20 Summarising the association of the aging phenotypes from the ROS/MAP study cohort and the ePRSs from $\Pi_{MHC/APOE-}$. The red dotted line represents 5% significance level. We outline the hub PRS of the highly associated module with the ageing phenotypes (smallest p-value, no module passed q-value $\leq$ 0.05 testing). These modules were selected to test if they can improve the predictability ability of state-of-the-art $PRS_{LOAD}$. . . . . . . . . . . . . . . . . . . . . . . . . . . . . . 30
S21 Summarizing the results of the multivariate association of modules from $\Pi_{MHC-}$. False discovery rate (FDR) correction (i.e. $q$-value [3]) was used to mitigate multiple testing concerns among all ePRSs (modules) on the likelihood ratio test between the full and base models. We selected ePRSs (modules) with q-value $\leq$ 0.05 and report it here. For continuous response variables (Beta Amyloid ($\beta$-amyloid) protein, PHF Tau, cognitive global random slope, cognitive global random slope at last visit) we report the .632 bootstrapped variation explained ($R^2$) by the covariates for the base (model 2) and full model (model 3). Additionally, we indicate the p-value of the likelihood ratio test between the full and base models. *$p \leq 0.05$, **$p \leq< 0.01$, ***$p \leq 0.001$. All p-values are reported at an uncorrected level. Note that the confidence intervals were measured out of the bootstrapping loop. 31

### 1. ANCESTRY BACKGROUND

We used a method incorporated in PLINK1.9 (www.cog-genomics.org/plink/): the multidimensional scaling (MDS) approach to account for population stratification. In general, the metric MDS, which is also known as the Principal Coordinates Analysis (PCoA), gives the same results as Principal Component Analysis (PCA) [4]. PCA is obtained by performing eigen-decompositions on a given matrix in order to achieve the eigen-configurations [5]. The given matrix (the similarity matrix) can be:

- Correlation matrix (variable are centered and normalized)
- Covariance matrix (variables are centered but not normalized)
- Cross product matrix (variables are neither centered nor normalized)

In order to calculate the similarity matrix of the metric MDS, Euclidean distance is used as the distance metric. All though a distance matrix cannot be analyzed directly using the eigen-decomposition (distance matrices are not positive semi-definite matrices), but they can be transformed into an equivalent cross-product matrix which can then be analyzed [5]. A metric MDS can be implemented by first defining an error function, as shown below:

$$S = \frac{\sum_{i,j} w_{ij}(d_{ij} - f(\delta_{ij}))^2}{\sum_{i,j} d_{ij}^2} \tag{S1}$$

where $w_{ij}$ are appropriately chosen weights, $\delta_{ij}$ is the dissimilarity between two points ($i, j$, which is the average proportion of alleles shared between any two individuals), and $d_{ij}$ is the interpoint distances. In metric MDS, we require

$$d_{ij} f(\delta_{ij}) \tag{S2}$$

where f is a specified function. By using gradient-based (or more sophisticated methods) to minimize the stress, one can obtain derivatives of $S$ based on the coordinates of the points that define $d'_{ij}s$ [4].

To investigate which individuals, the generated component scores deviate from the sample target population, plotting the sample scores under investigation and a population of known ethnic structure (e.g., ROS/MAP vs 1000 Genome data) is helpful: This step is called anchoring [6]. Quantitative components of genetic variations were generated for each individual based on the average proportion of alleles shared between any two individuals within a sample. If the individual component scores of the individuals are plotted against each other, it can be explored whether or not there are any groups of genetically similar individuals that did not occur as expected. The MDS plots of the ROS/MAP genotype is shown in Figure S2. They verify the homogeneity (Caucasian European) of the population of the study. Additionally, to account for stratification in our study, we used the first 10 PCs in our analysis.

### 2. GWAS SUMMARY STATISTICS

For phenome-wide PRS calculations, we used GWAS summary statistics derived from the UK Biobank, a large-scale population-based study including approximately 500,000 individuals [7]. The Pan-UK Biobank consortium has recently conducted GWAS (www.pan.ukbb.broadinstitute.org) for all phenotypes with sufficient statistical power. As described in www.pan.ukbb.broadinstitute.org, each GWAS conducted for each phenotype (for each ancestry group) was used from imputed variants from the UK Biobank, with 97,059,328 variants. Then the variants with INFO score 0.8 were retained (29,865,259 variants on the autosomes and X-chromosome). Additionally, variants with allele count of at least 20 were filtered. The model used for each GWAS is shown below:

$$Y_{Phenotype} \sim Age + Sex + Age * Sex + Age^2 + Age^2 * Sex + \sum_{i=1}^{2} PC_i \tag{S3}$$

The heritability scale and genomic control ($\lambda_{GC}$) for each phenotype was measured and reported with each GWAS. These measurements were used to verify the quality of each GWAS summary statistics. Genomic control (GC) is used to correct $\chi^2$ test statistics which are assumed to be inflated by a factor $\lambda_{GC}$. [8]. It is defined as the median of the resulting $\chi^2$ test statistics divided by the expected median of the $\chi^2$ distribution [8].

In each GWAS summary statistics, the population stratification was corrected by the first 10 PCs. As shown in previous work by [9] the PCA correction method can control for type I error well, and it has a much higher power in meta-analysis compared to the GC correction method, where the allele effect is divided by $\lambda_{GC}$. Additionally, other studies have shown in polygenic inheritance, substantial genomic inflation is expected, and its magnitude depends on sample size, heritability, linkage disequilibrium structure and the number of causal variants [10].

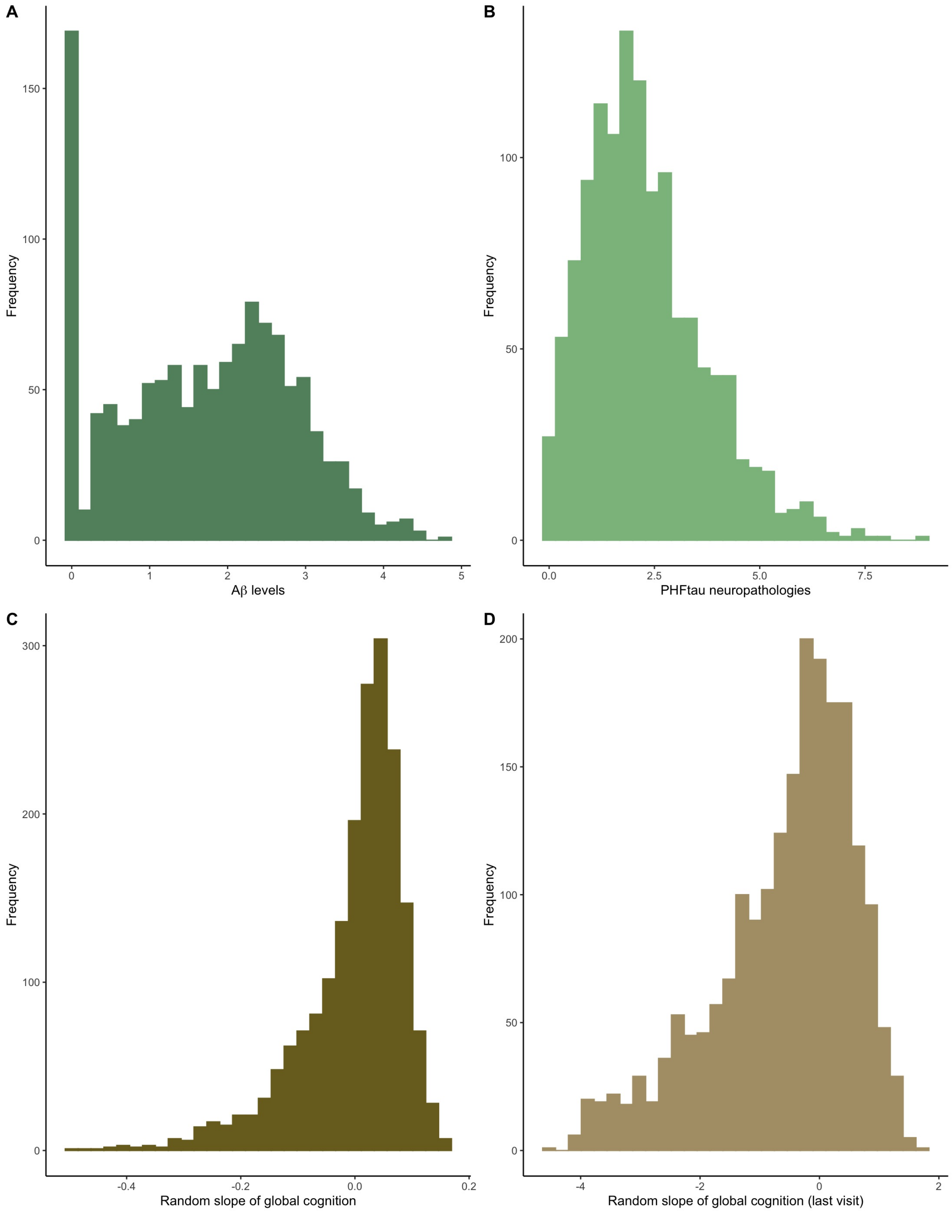

Fig. S1: Distribution of postmortem pathology and rates of antemortem cognitive decline calculated in n=2,044 participants from the Religious Orders Study and Memory and Aging Project (ROS/MAP). **A**. $\beta$-Amyloid protein (n=1,238) identified by molecularly-specific immunohistochemistry and quantified by image analysis. Value is percent area of cortex oocupied by amyloid beta. Mean of $\beta$-Amyloid score in 8 regions [2]. **B**. Neuronal neurofibrillary tangles (n=1,247) are identified by molecularly specific immunohistochemistry (antibodies to abnormally phosphorylated Tau protein, AT8). Cortical density is determined using systematic sampling. Mean of tangle score in 8 regions (4 or more regions are needed to calculate) [2]. **C**. Measurement of global cognitive decline (n=1,915). **D**. Last measurement of global cognitive decline (n=2,040).

### A. Selection of GWAS summary statistics

Out of a total of 7,221 phenotypes with GWAS performed in the European ancestry UKB sample (which matches our sample of genetically verified European-ancestry ROS/MAP participants), we selected 2,218 phenotypes that had heritability estimates greater than or equal to 5% and were sex non-specific. The heritability scale of each phenotype was provided by Pan-UKBB consortium (calculated using SAIGE [11]), and selected sex non-specific phenotypes using their provided meta-data including some additional manual selection; e.g. country of birth, age at menopause, blood pressure measurement using different methods (manual or electronic, right or left hand). Sex non-specificity was used as a criterion to ensure that all PRS were meaningful for all ROS/MAP participants (e.g. PRS for vasectomy is not meaningful for female participants). As a benchmark for PRS predictive performance of LOAD, we also accessed the latest summary statistics of LOAD [1], which represent the current standard.

Here, we outline a list of selection criteria we applied to the GWAS summary statistics:

- Any phenotype with hertiablity ratio of less than 5% were removed (4,496 phenotypes were removed).
- Inclusion of both sexes in the cases for each phenotype. This was done through the `pheno_sex` variable provided by the PANUKBB consortium (44 phenotypes were removed).
- Removing any non European population in the GWASs (none of the phenotypes were removed).
- Some of the phenotypes were related to country of birth. We removed such phenotypes to avoid any potential stratification bias in our analysis (35 phenotypes were removed).
- Some phenotypes were predominantly sex-specific; However, both sexes can develop such conditions like breast cancer. Thus, we removed such phenotypes from the summary statistics (16 phenotypes were removed).
- We used:
  1. `n_cases_full_cohort_females`
  2. `n_cases_full_cohort_males`

  to make sure there were cases of both male and female in the GWAS results for each phenotype (230 phenotypes were removed).
- Any administrative phenotypes (e.g. location of the hospital patient was registered at, or location of patient's home to the hospital) were manually removed (118 phenotypes were removed).
- Phenotypes that are repeated measurement of the same condition. For example, pulse pressure measured manually or with a digital device. Thus, we included one of such phenotypes by either selecting the one with ICD coding, or one with the highest heritability ratio (62 phenotypes were removed).
- Two LOAD summary statistics from the PANKUKBB summary statistics were removed. Since we used a different GWAS summary statistics for LOAD.

An external table is provided to outline the list of phenotypes selected in this study.

(`Table_S3_ukbb_manifest_filtered_phenotypes.csv`)

Table S1: In this table we outline the selected GWAS summary statistics of phenotypes used in this study. The GWAS summary statistics were taken from the Pan-UK biobank consortium, except for GWAS of LOAD which was taken from [1]. Each column is previously defined by the Pan-UK biobank (except for LOAD). Phenocode is the code used for the phenotypes, and trait type is one of the following: continuous, biomarkers, prescriptions, icd10, phecode, categorical. The phenotype name is the name of each phenotype as they appear in this study. ICD10 is the coding of the phenotypes defined by the International Classification of Diseases. Cases and control represent the number of case and control (only European population) in each GWA study. Heritablity is the saige heritablity measure of each GWAS (except for LOAD). $\lambda_{GC}$ is the genomic control of each GWAS (except for LOAD). Lastly the last column provides the link to each summary statistics.

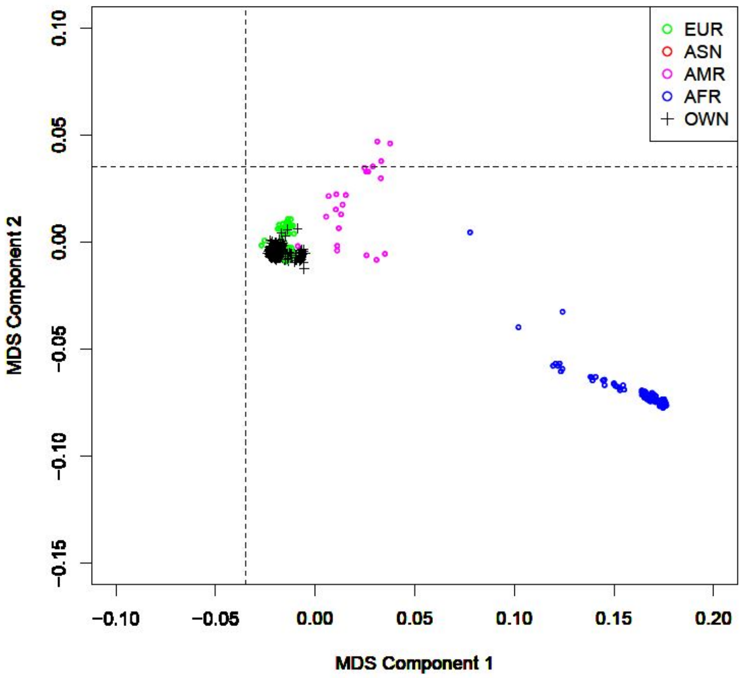

((a))

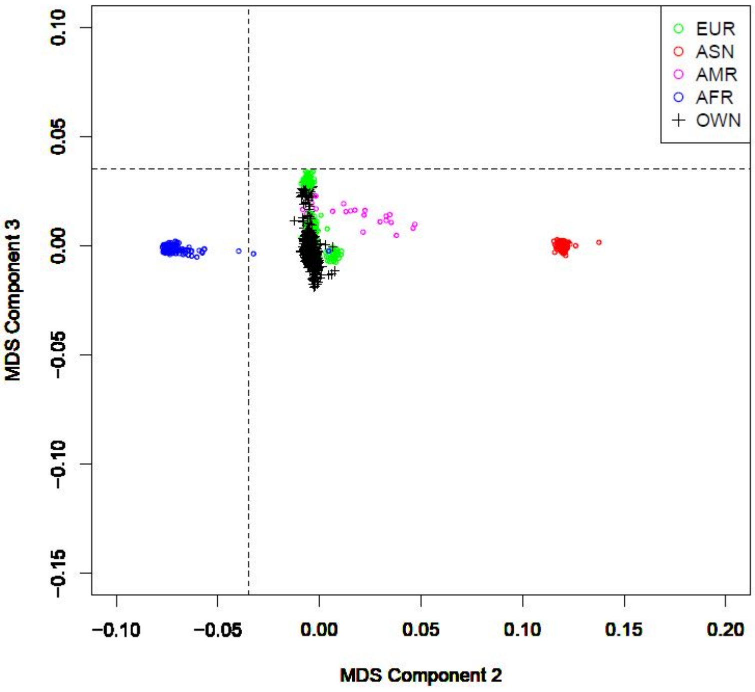

((b))

Fig. S2: Multidimensional scaling (MDS) plot of 1000 Genome against the ROS/MAP dataset 2,067 individuals. The black crosses (+='OWN') in the upper left part represent the first two MDS components of the individuals in the ROS/MAP dataset (The colored symbols represent the 1000 Genome data ('EUR'=European; 'AFR'=African; 'AMR'=Ad Mixed American; 'ASN'=Asian)). The MDS components representing the European samples (the green color) is overlapped with our dataset, showing the homogeneity of our population. Each figure S2(a,b) are showing the MDS plot in different dimensions.

| | $\Pi_{ALL}$ | | $\Pi_{MHC-}$ | | $\Pi_{APOE-}$ | | $\Pi_{MHC/APOE-}$ | |
|---|---|---|---|---|---|---|---|---|
| $\alpha$-value | before QC | after QC | before QC | after QC | before QC | after QC | before QC | after QC |
| 1 | 2218 | 1967 | 2218 | 1967 | 2218 | 1965 | 2218 | 1967 |
| 0.1 | 2218 | 2012 | 2218 | 2012 | 2218 | 2010 | 2218 | 2011 |
| 0.05 | 2218 | 2027 | 2218 | 2030 | 2218 | 2025 | 2218 | 2030 |
| 0.01 | 2218 | 2058 | 2218 | 2062 | 2218 | 2055 | 2218 | 2064 |
| 0.005 | 2218 | 2064 | 2218 | 2068 | 2218 | 2064 | 2218 | 2068 |
| 0.001 | 2218 | 2065 | 2218 | 2074 | 2218 | 2065 | 2218 | 2074 |
| 0.0005 | 2218 | 2070 | 2218 | 2075 | 2218 | 2068 | 2218 | 2075 |
| 0.0001 | 2218 | 2060 | 2218 | 2070 | 2218 | 2062 | 2218 | 2073 |
| 5.00e-05 | 2218 | 2060 | 2218 | 2072 | 2218 | 2061 | 2218 | 2074 |
| 1.00e-05 | 2215 | 2043 | 2214 | 2065 | 2215 | 2039 | 2215 | 2065 |
| 5.00e-06 | 2027 | 1843 | 2008 | 1848 | 2030 | 1843 | 2007 | 1848 |
| 1.00e-06 | 779 | 631 | 705 | 594 | 791 | 644 | 703 | 593 |
| 5.00e-07 | 642 | 511 | 566 | 465 | 648 | 516 | 560 | 461 |
| 1.00e-07 | 520 | 395 | 452 | 363 | 517 | 396 | 443 | 353 |
| 5.00e-08 | 476 | 349 | 412 | 321 | 475 | 350 | 405 | 315 |

Table S2: Total number of PRSs before (before QC) and after (after QC) applying initial quality control. At each $\alpha$-value threshold (first column), we removed PRSs which were calculated with less than 5 SNPs. We also removed duplicated PRSs, as described in methodology. We did this process across all PRS matrices ($\Pi_{ALL,\alpha}$, $\Pi_{MHC-,\alpha}$, $\Pi_{APOE-,\alpha}$, $\Pi_{MHC/APOE-,\alpha}$).

### 3. PRINCIPAL COMPONENT ANALYSIS (PCA)

PCA, a multivariate technique that can describe the inter-correlations amongst the dependent variables (PRSs of up to 2,075 phenotypes) within the matrix [12]. Theoretically, the principal components of the PRSs uncovers the dominant combinations of phenotypes (genetics) that describe as much of the PRS matrix as possible. The goals of PCA are to:

1. Identify the most important information in the original data.
2. Keep the most important information in the data set and reduce the data set size.
3. Provide a simplified description of the data set.
4. Analyze the structure of the observations and the data variables.

Algebraically, PCA is the eigendecomposition of the covariance matrix of the original matrix $\Pi_{N\times K}$. The components are obtained from the singular value decomposition (SVD) of $\Pi$, specifically with:

$$\Pi = P\Sigma = U\Sigma V^T \tag{S4}$$

Where $P_{N\times L}$ is the matrix of factor scores, and $V_{K\times L, L\leq min\{N,K\}}$ (in our case the components or the PRSs) is the coefficients of the linear combinations used to compute the score matrix $P_{N\times K}$ (factors scores or projection matrix). It is denoted as:

$$P = U\Sigma = U\Sigma V^T V = \Pi V \tag{S5}$$

#### A. PRS Contributing Score

An observation's importance for a component can be determined by the ratio between its squared factor score ($F_{n\times l}$) and its eigenvalue. This ratio is called the contribution of the observation to the component. The contribution of observation $n$ to component $l$, denoted by $ctr_[n, l]$, can be expressed as follows:

$$ctr_{n,l} = \frac{p_{n,l}^2}{\sum_i f_{n,l}^2} \tag{S6}$$

The contributing score for a given PRSs to a given PC is between 0 and 1, and for a given component, the sum of contributions of all observations is equal to 1. It is, therefore, possible to quantify them as a percentage. An observation is considered to be contributing more to a component if its contribution value has a higher value [12].

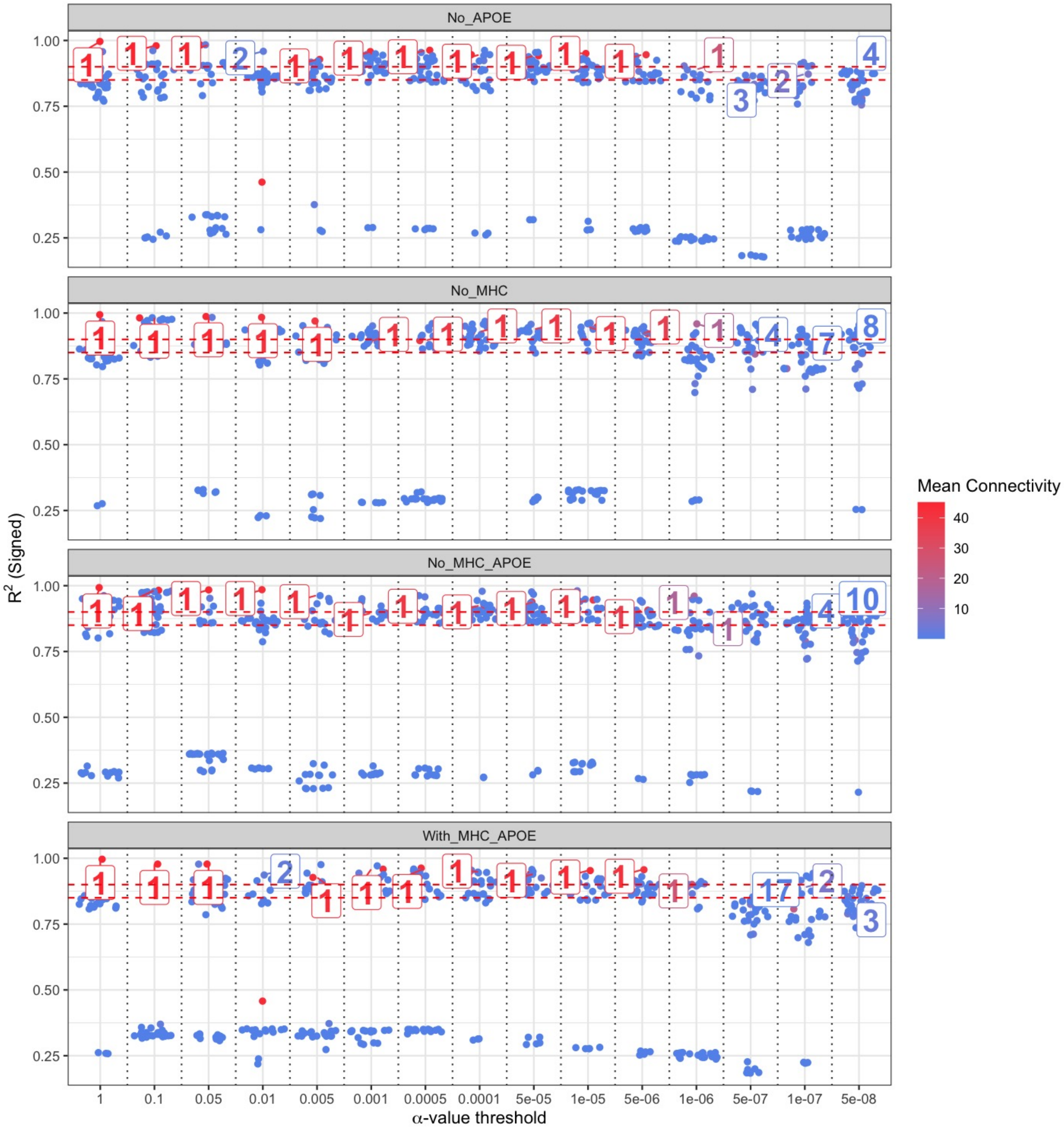

Fig. S3: To achieve Scale-Free Topology, we select the smallest $\beta$ value so that the $R^2$ of equation S14 is highest. The $\beta$ values selected in constructing the Scale-Free Topology are labelled in this plot. As $\beta$ increases, we have smaller mean connectivity (refer to equation S13). Additionally, we notice that with higher $\alpha$-value threshold, $\beta = 1$, indicating that by including every gene in our PRS calculations for all phenotypes and the 2,044 ROS/MAP participants, we obtain a scale-free topology.

### 4. WEIGHTED GENE CO-EXPRESSION NETWORK ANALYSIS (WGCNA)

This section will discuss how we constructed our network and how we were able to detect the PRS modules in the network.

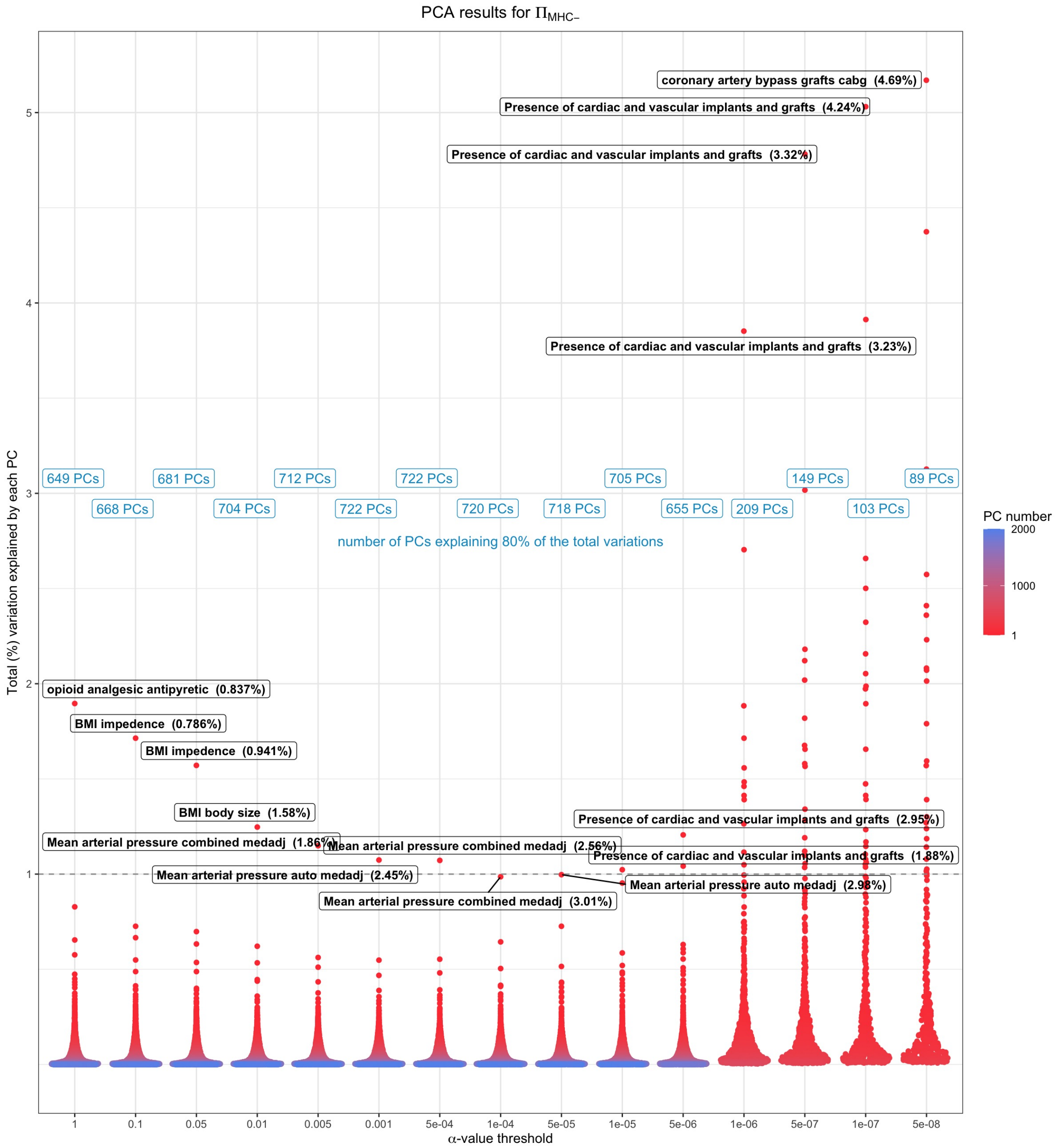

Fig. S4: Principal Component Analysis of phenome-wide PRS. This plot outlines the results from the PC analysis on $\Pi_{MHC-}$. Each point is a component for each matrix, and the colour of each point represents the PC number. The top contributor of each PC, with their contributing percentage to the component is shown for the first PC of each matrix. These PCs are mainly dominated by autoimmune diseases at lower $\alpha$-value thresholds, and general pain medications at higher $\alpha$-value thresholds.

### A. Adjacency Matrix

The adjacency matrix $a_{ij}$ is a symmetric matrix with entries in [0, 1] whose component $a_{ij}$ represents the degree of network connection strength between nodes i and j, representing a symmetric N by N matrix. It is the adjacency matrix of a network that fully specifies the nature of the network. The adjacency matrix is calculated by the

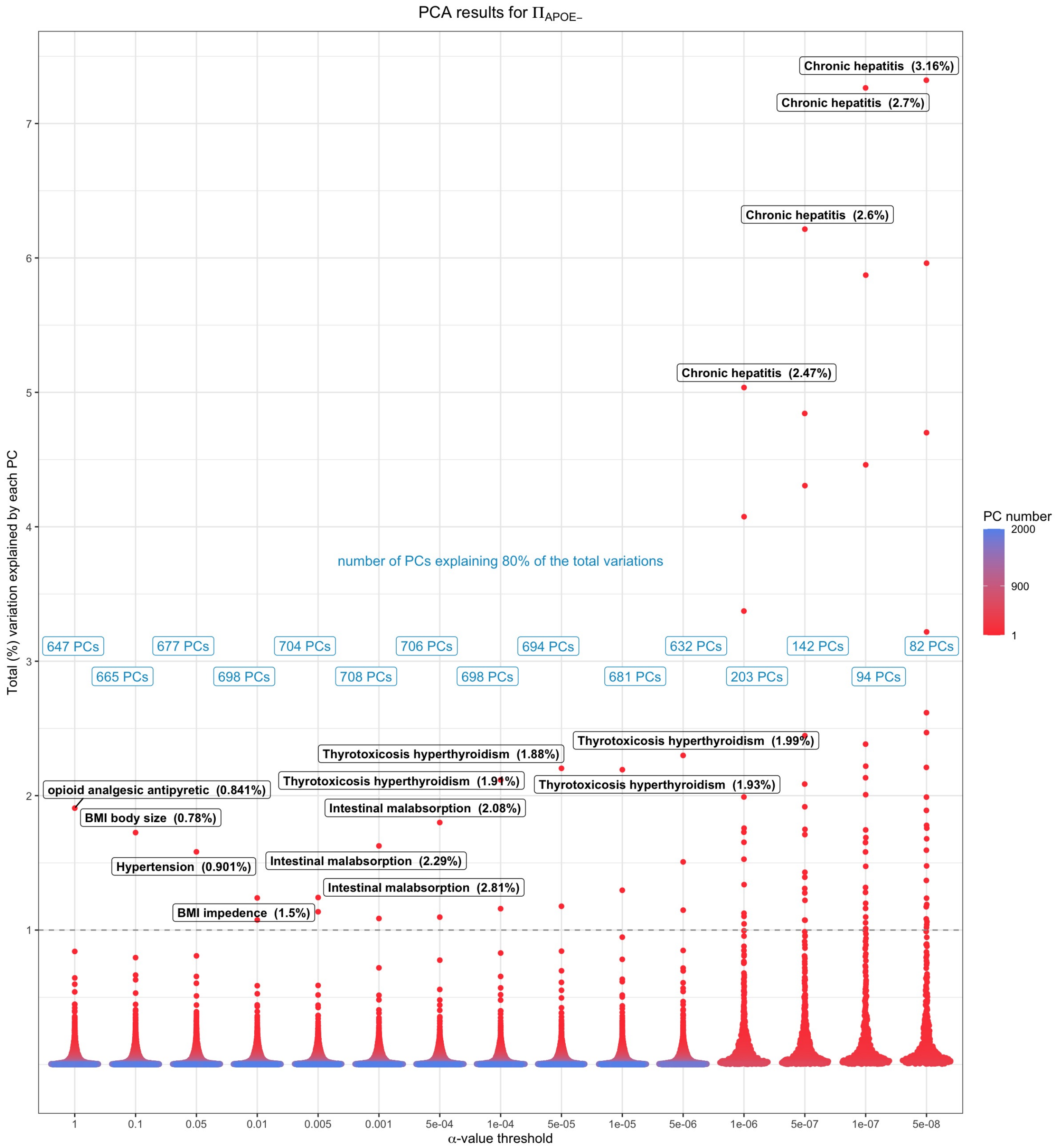

Fig. S5: Principal Component Analysis of phenome-wide PRS. This plot outlines the results from the PC analysis on $\Pi_{APOE-}$. Each point is a component for each matrix, and the colour of each point represents the PC number. The top contributor of each PC, with their contributing percentage to the component is shown for the first PC of each matrix. These PCs are mainly dominated by autoimmune diseases at lower $\alpha$-value thresholds, and general pain medications at higher $\alpha$-value thresholds.

intermediate quantity, called the co-expression similarity $s_{ij}$. As a default method, the co-expression similarity $s_{ij}$ is defined to be the absolute value of the correlation coefficient amongst the profiles of nodes i and j: $s_{ij} = |cor(x_i, x_j)|$. Pearson correlation was used to evaluate the similarities between the PRSs.

A thresholding procedure is used in order to transform the PRS similarity (originally reffered to co-expression similarity) into the adjacency. An unweighted network adjacency $a_{ij}$ between PRS profiles $x_i$ and $x_j$ can be defined

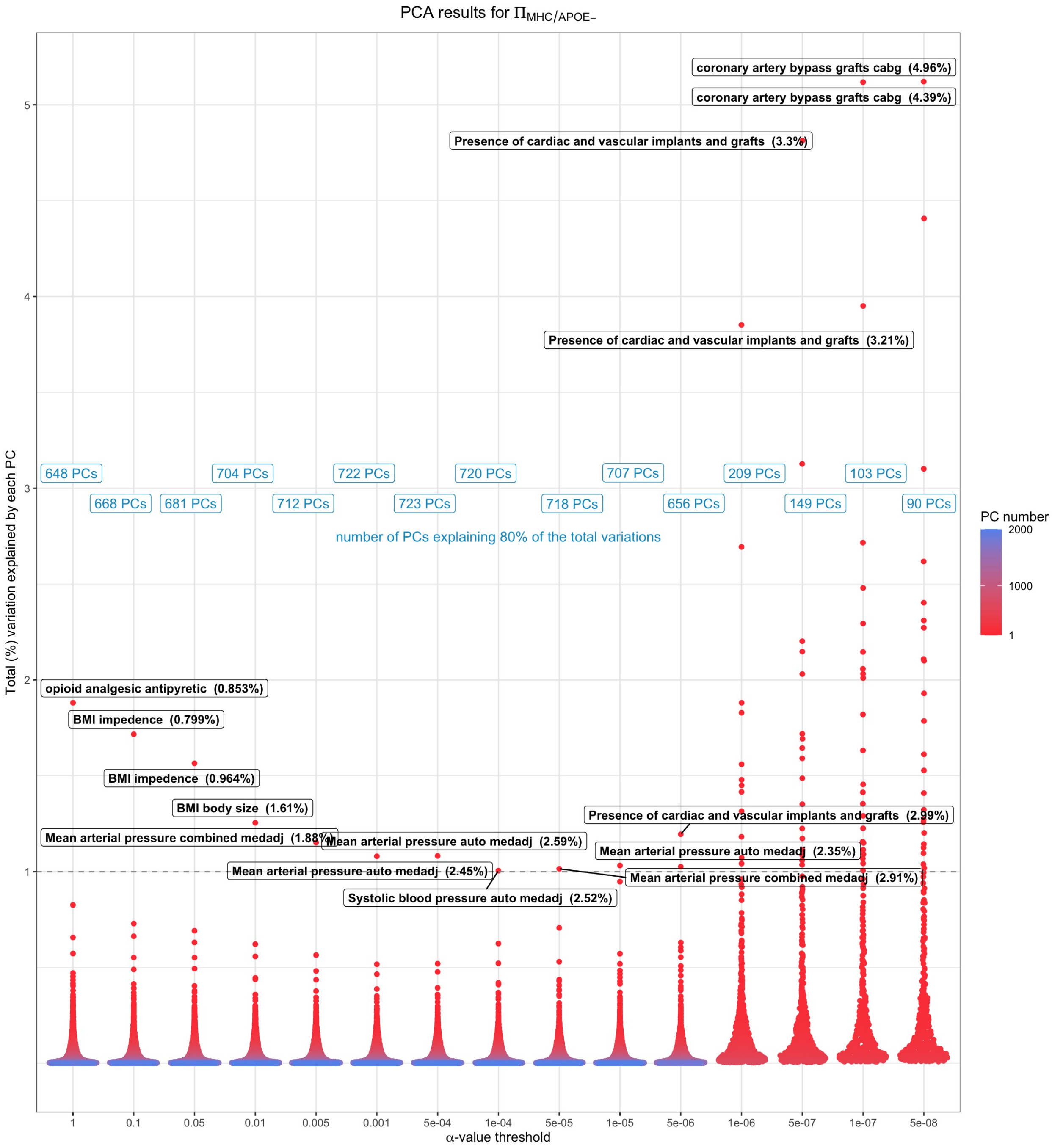

Fig. S6: Principal Component Analysis of phenome-wide PRS.This plot outlines the results from the PC analysis on $\Pi_{MHC/APOE}$. Each point is a component for each matrix, and the colour of each point represents the PC number. The top contributor of each PC, with their contributing percentage to the component is shown for the first PC of each matrix. These PCs are mainly dominated by autoimmune diseases at lower $\alpha$-value thresholds, and general pain medications at higher $\alpha$-value thresholds.

by hard thresholding the PRS similarity $s_{ij}$ as

$$a_{ij} = 1 s_{ij} \geq \tau 0 otherwise \tag{S7}$$

where $\tau$ is the "hard" threshold parameter. While unweighted netwroks are widely used, they do not reflect the continuous nature of the PRSs. In the context of weighted networks, the adjacency can have values which range from 0 to 1. The PRS similarity can be raised to a power ($\beta \geq 1.$) to define a weighed network adjacency [13] [14].

$$a_{ij} = s_{ij}^{\beta} \tag{S8}$$

The adjacency in Equation 3.6 implies that the weighted adjacency $a_{ij}$ between two PRSs is proportional to their similarity on a logarithmic scale, $log(a_{ij}) = \beta \times log(s_{ij})$. The threshold parameters are selected to obtain a scale free topology. Based on the continuous nature of the PRSs, we used the soft thresholding method on our weighted network.

In the weighted networks, $a_{ij}$ is measured in two methods:

1. Unsigned network:
$$a_{ij} = \|cor(x_i, x_j)\|^{\beta} \tag{S9}$$
2. Signed network (preserves the sign):
$$a_{ij} = \|0.5 + 0.5cor(x_i, x_j)\|^{\beta} \tag{S10}$$

where $\beta$ is the soft thresholding and $x_i$ and $x_j$ are the column vectors from the $\Pi_{nk}$ matrix. Each vector would represent $\Pi_{subscript}$ of one phenotype for all ROS/MAP participants. Thus the adjacency matrix of the PRSs will be a k by k matrix, representing the similarities between every possible combination of the PRS vectors. We calculated an unsigned network, meaning that adjacency was based on the absolute value of correlation rather than discarding negative correlations. This was important as we calculated PRS across thousands of phenotypes, among which both negative and positive correlations would interest us. In our analysis, for each PRS matrix, pairwise Pearson correlations were calculated among all PRSs.

### B. A Scale-Free Network

For an undirected (our network is undirected, since we are interested in correlation between the PRSs) network, we can write the degree distribution as:

$$P_{deg}(k) \sim k^{-\gamma} \tag{S11}$$

,where $\gamma$ is some exponent whose value is typically in the range $(2, 3)$, however it may lie outside of these range occasionally [15] [16] [17]. $P_{deg}(k)$ decays slowly as the degree k increases, increasing the likelihood of finding a node with a very large degree. Additionally k is calculated as

$$k_i = \sum_j a_{ij} \tag{S12}$$

,which is the row sum of the adjacency matrix. For a network to satisfy a scale-free topology, they are highly dependent on the $\beta$ threshold. Thus, we choose the power, $\beta$ so our network satisfies a scale-free topology. Generalizing the notion of scale-free topology from previous studies we have:

$$log(P(k_{deg})) = b_0 + b_1 log(k)[15] \tag{S13}$$

We choose the $\beta$ parameter so the model in equation S14 has the best goodness of fit ($R^2$ close to 1). For the construction of a weighted network, the PRS-PRS correlations were raised to a soft-thresholding power ($\beta$) via the WGCNA package function pickSoftThreshold.

### C. Module Detection

Once the network is fully constructed, WGCNA detects modules of PRSs based on their similarities. Modules are defined as clusters of densely interconnected PRSs. Several ways of measuring network interconnectedness are described in [18]. By default, WGCNA uses the topological overlap measure, since it has worked in several applications [13] [19] [20]. The topological overlap of two nodes relates to their similarity with respect to the commonality of nodes they are connected to, as reflected in [18].

#### D. Calculations of Topological Properties

The topological overlap of two nodes reflects their similarity in terms of the commonality of the nodes they connect to [18]. [21] and [22] define the topological overlap matrix $T = [t_{ij}]$ as follows:

$$t_{ij} = \frac{l_{ij} + a_{ij}}{min(k_i, k_j) + 1 - a_{ij}}, i \neq j, i = j \tag{S14}$$

Where, $l_{ij} = \sum_u a_{iu}a_{uj}$, $k_i = \sum_u a_{iu}$ and the index $u$ runs across all nodes of the network. Additionaly, $k_i$ is the sum of $i^{th}$ row of the adjacency matrix, and $k_j$ is the sum of $j^{th}$ column of the adjacency matrix. Basically, $t_{ij}$ is an indicator for the agreement between the sets of neighbouring nodes of $i$ and $j$. The inclusion of the term $a_{ij}$ in the numerator makes $t_{ij}$ explicitly depends on whether there is a direct link between the two nodes in question. The purpose of the quantity $1 - a_{ij}$ in the denominator is to avoid double-counting $i$ as a neighbor of $j$ and vice versa [22].

### 5. CLUSTERING OF PRS MATRICES AND CALCULATION OF EIGEN-PRSS (EPRSS) USING WGCNA

For each PRS matrix, the topological overlap matrix (TOM) was calculated and used to define the hierarchical clustering dissimilarities. The resulting dendrogram was then clustered using Dynamic Tree Cutting [23] which applies a variable-height tree cut algorithm to maximize the capture of meaningful branch structures (i.e. PRS modules). WGCNA was run using the following additional parameters for detection of PRS modules: module detection cut-height of 0.999, minimum module size of 10, and high branch split sensitivity (deepSplit=4).

Our soft-thresholding power selection varied across PRS matrices, mainly depending on the SNP inclusion $\alpha$-value threshold and, therefore, the number of PRSs included in a given matrix. For each matrix, the lowest value of $\beta$ was chosen at which the scale-free topology model fit ($R^2$) reached above 0.8 (resulting $\beta$ values ranging from 1 to 14) to satisfy a scale-free topological network. In addition, we set a very low minKMEtoStay parameter (0.01), which governs the behaviour of the dendrogram clustering algorithm and makes nearly all input PRSs members of at least one module (instead of excluding lowly correlated PRSs from membership in any module).

Following the PRS matrix clustering and the identification of PRS modules, PCA was applied to each module separately, with the first principal component representing that module's eigen-PRS (ePRS); multiple principal components can be used for each module, but most of the ePRSs capture over 40% of the variation for modules at lower $\alpha$-value thresholds; thus used to simplify the downstream analysis. These ePRSs were then 1) examined for insights into the whole-person genetic risk landscape across input PRS matrices and 2) used for downstream modelling of LOAD phenotypes.

#### A. Module Preservation

In order to grasp a better understating of the modules generated by WGCNA, we looked into the most preserved modules. This allowed us to check the robustness of the modules produced by WGCNA at different $\alpha$-value thresholds. Statistically, the network-based module preservation can be tested by different methods [24]. We used connectivity to test (If the hub status preserved between reference and test networks?) for module preservation. First, we identified the hub PRSs preserved at every $\alpha$-value threshold. By measuring the pairwise correlation of their corresponding ePRS, we identify the correlation structure of the hub PRS within each reference ($\Pi_{ALL}$) and test ($\Pi_{MHC-}$, $\Pi_{APOE-}$, $\Pi_{MHC/APOE-}$) network.

Figures S12 to S15 outline the mostly connected PRSs (hub PRSs) for each module and the number of times they were repeated at every $\alpha$-value threshold. We removed any module that appeared once across all thresholds (except for the modules represented by family history of AD dementia). We also chose a very low minKMEtoStay parameter (0.01), which governs the behaviour of the dendrogram clustering algorithm and results in the inclusion of nearly all input PRS into some module (rather than excluding lowly correlated PRSs from membership in any module). Hub PRS represented by weight was preserved at every $\alpha$-value threshold in $\Pi_{ALL}$ except for 0.1 and 5e-08, it was also preserved in $\Pi_{APOE-}$ except at $\alpha$-value threshold of 5e-08.

Additionally, the hub PRS for high cholesterol, which was highly associated with LOAD, was mainly preserved at lower $\alpha$-value thresholds. However, the high cholesterol hub did not appear to be preserved when the *APOE* region was excluded ($\Pi_{APOE-}$, $\Pi_{MHC/APOE-}$). Interestingly, with the removal of the MHC region, we noted hub PRS of asthma was the most preserved module across all $\alpha$-value thresholds (except for $\alpha$ threshold of 1, 5e-06, 1e-07, and 5e-08). It is important to note that not all PRS memberships is a preserved module are the same at different $\alpha$-value thresholds. Thus, we seeked to identify the correlation structure of some of the selected modules (e.g. weight, high cholesterol) to verify the module preservation and robustness. The heatmaps outlined in Figure S16, verify the robustness of both weight and high cholesterol hubs. Even with the removal of the *APOE* region or the MHC region, we notice that the hub PRSs of weight are highly correlated with each other.

| $\alpha$-value threshold | $\Pi_{ALL}$ | $\Pi_{MHC-}$ | $\Pi_{APOE-}$ | $\Pi_{MHC/APOE-}$ |
|---|---|---|---|---|
| 1 | 35 | 33 | 34 | 34 |
| 0.1 | 38 | 35 | 37 | 35 |
| 0.05 | 33 | 32 | 33 | 32 |
| 0.01 | 61 | 23 | 62 | 24 |
| 0.005 | 16 | 16 | 16 | 15 |
| 0.001 | 10 | 15 | 10 | 15 |
| 0.0005 | 10 | 14 | 10 | 14 |
| 0.0001 | 11 | 18 | 11 | 18 |
| 5.00e-05 | 13 | 19 | 13 | 20 |
| 1.00e-05 | 16 | 28 | 16 | 25 |
| 5.00e-06 | 15 | 27 | 16 | 27 |
| 1.00e-06 | 15 | 18 | 15 | 18 |
| 5.00e-07 | 8 | 14 | 14 | 15 |
| 1.00e-07 | 13 | 10 | 12 | 10 |
| 5.00e-08 | 12 | 8 | 12 | 7 |

Table S3: Here you will find a list of the number of modules selected by the WGCNA based on all $\alpha$-value thresholds and genome region inclusion criteria. The highest number of modules was 62, constructed from $\Pi_{APOE-,\alpha\leq0.01}$.

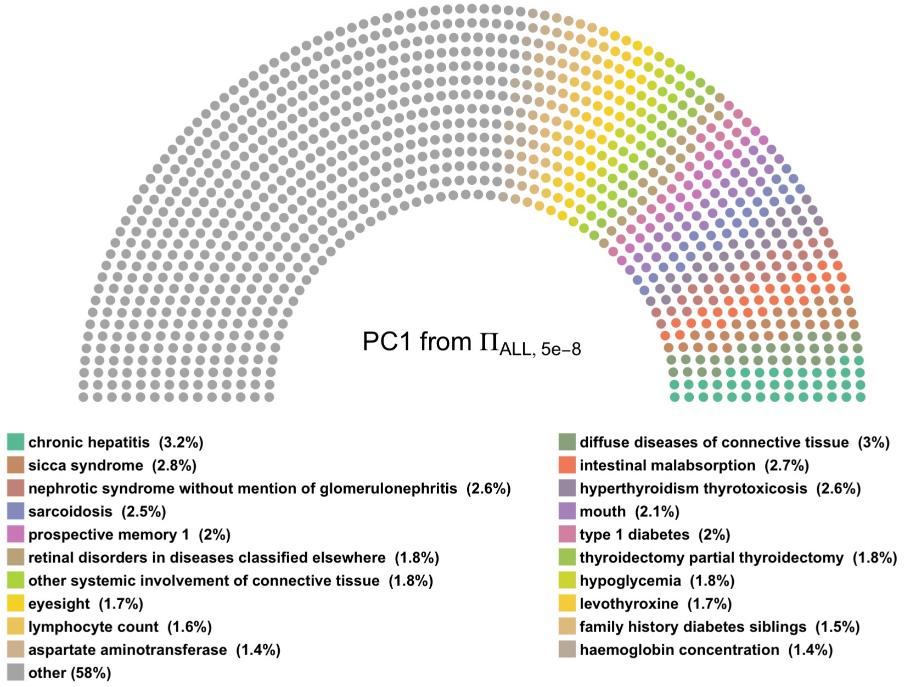

**((c))** This PC explained 7.38% of the total variation of the PRS matrix.

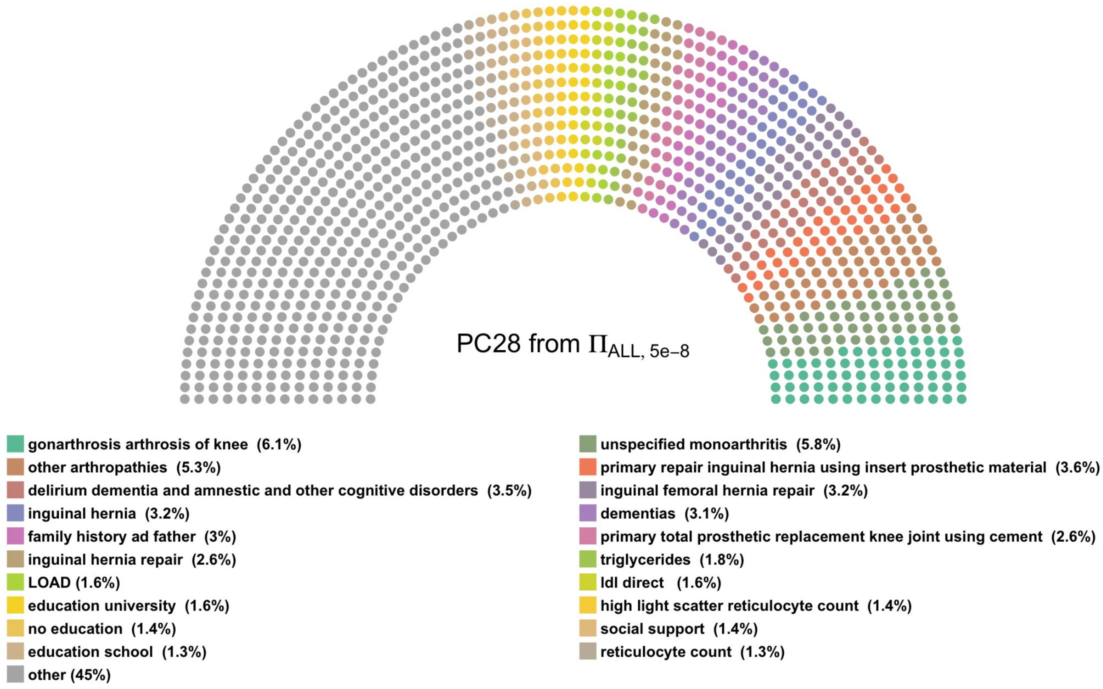

**((d))** This PC explained 0.83% of the total variation of the PRS matrix.

Fig. S7: Top 25 contributors to PC1 and PC28, calculated from $\Pi_{ALL}$, are shown with their contribution. Each dot represents 0.4% of the PRS contribution to this PC.

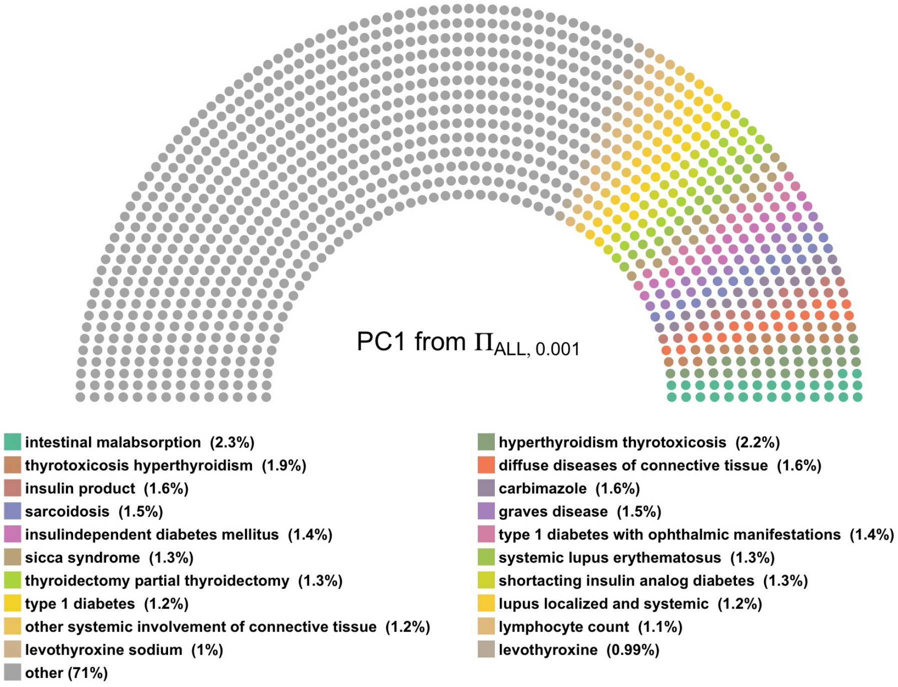

**((e))** This PC explained 1.63% of the total variation of the PRS matrix.

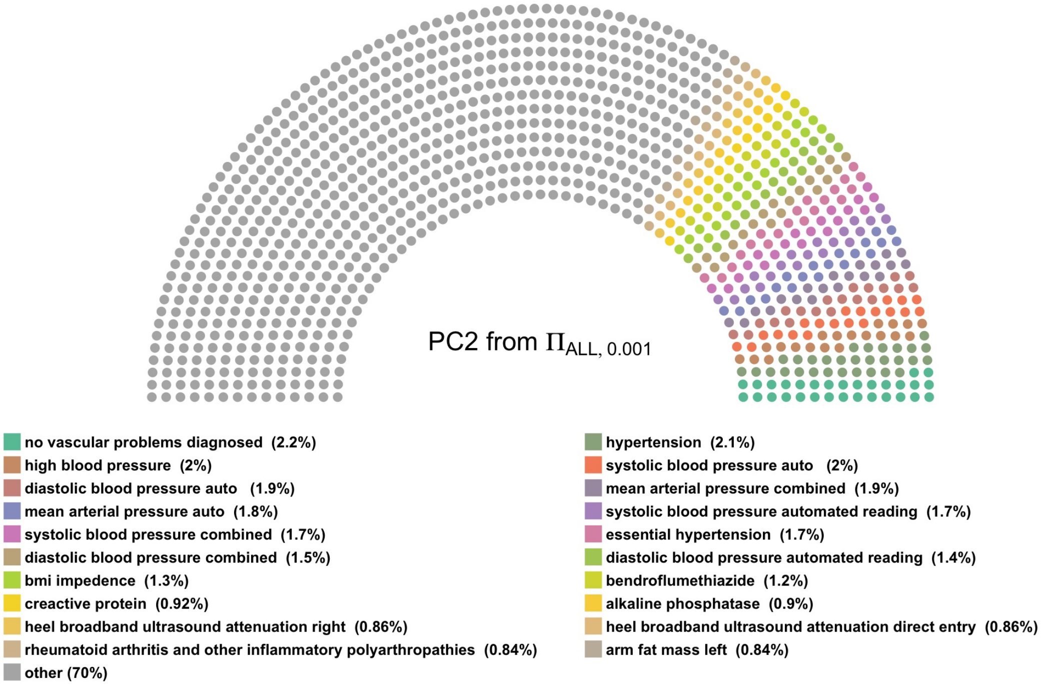

**((f))** This PC explained 1.06% of the total variation of the PRS matrix.

Fig. S8: Top 25 contributors, with their contribution, to PC1 and PC2, calculated from $\Pi_{ALL,0.001}$, are shown here. Each dot represents 0.4% of the PRS contribution to this PC.

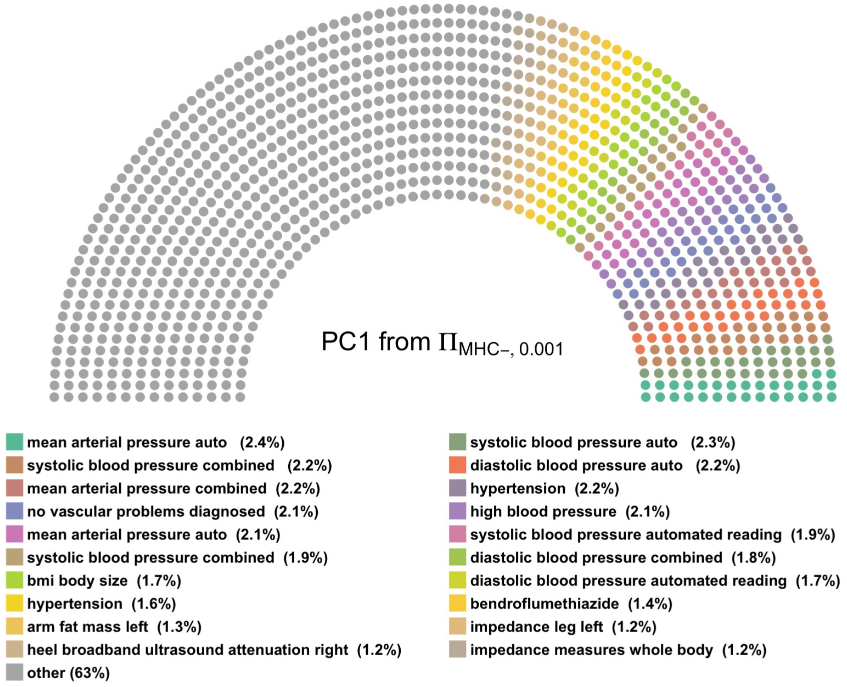

**((g))** This PC explained 1.07% of the total variation of the PRS matrix.

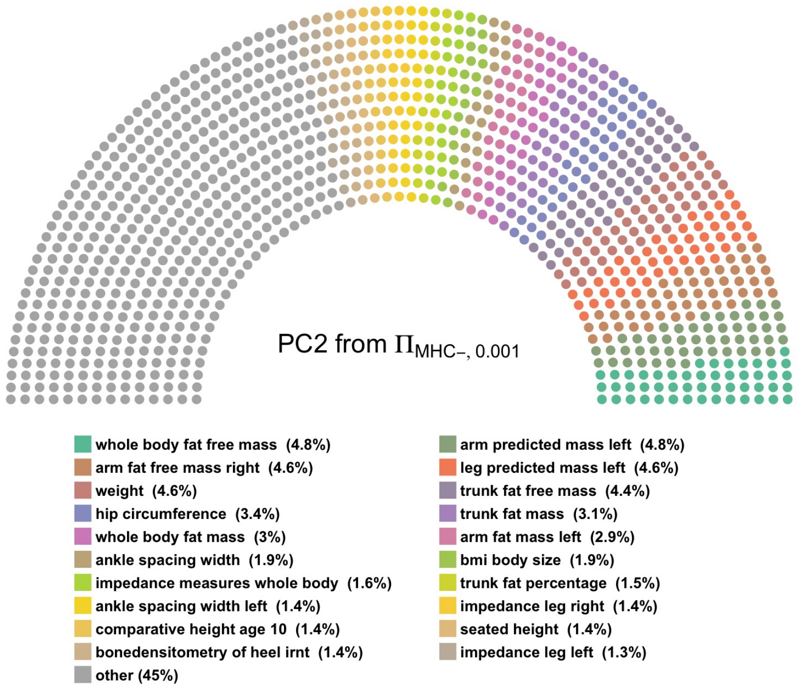

**((h))** This PC explained 0.55% of the total variation of the PRS matrix.

Fig. S9: Top 25 contributors, with their contribution, to PC1 and PC2, calculated from $\Pi_{MHC-,0.001}$, are shown here. Each dot represents 0.4% of the PRS contribution to this PC.

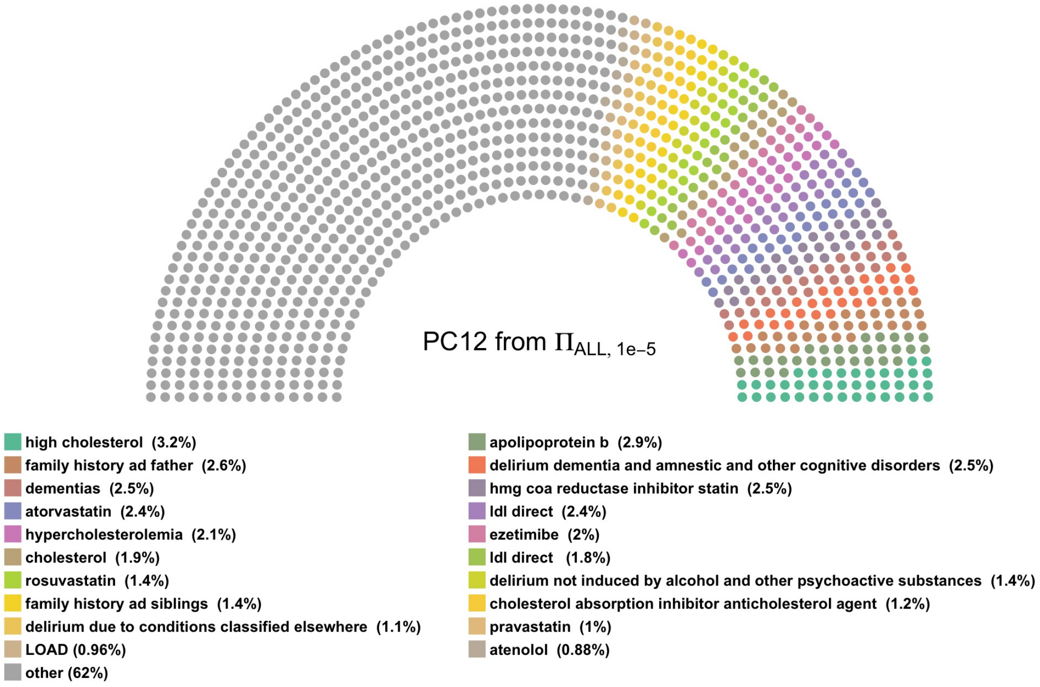

**((i))** This PC explained 0.37% of the total variation of the PRS matrix.

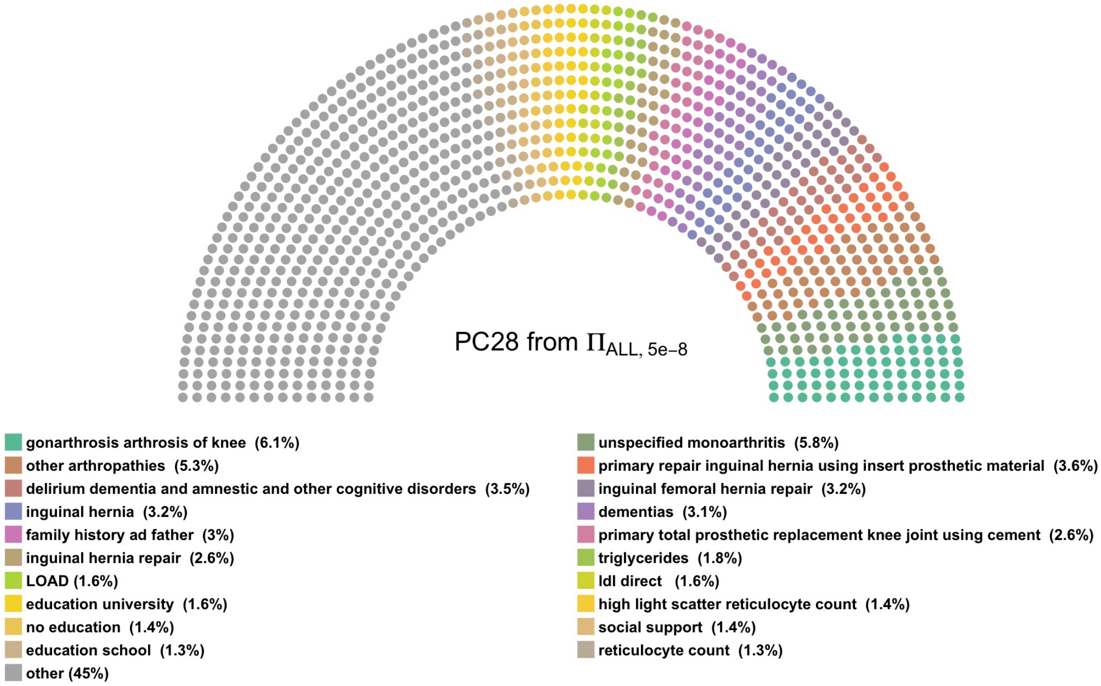

**((j))** This PC explained 0.83% of the total variation of the PRS matrix.

Fig. S10: Top 25 contributors, with their contribution, to PC12 and PC9, calculated from $\Pi_{ALL,1e-05}$ and $\Pi_{ALL,5e-08}$, are shown here. Each dot represents 0.4% of the PRS contribution to this PC.

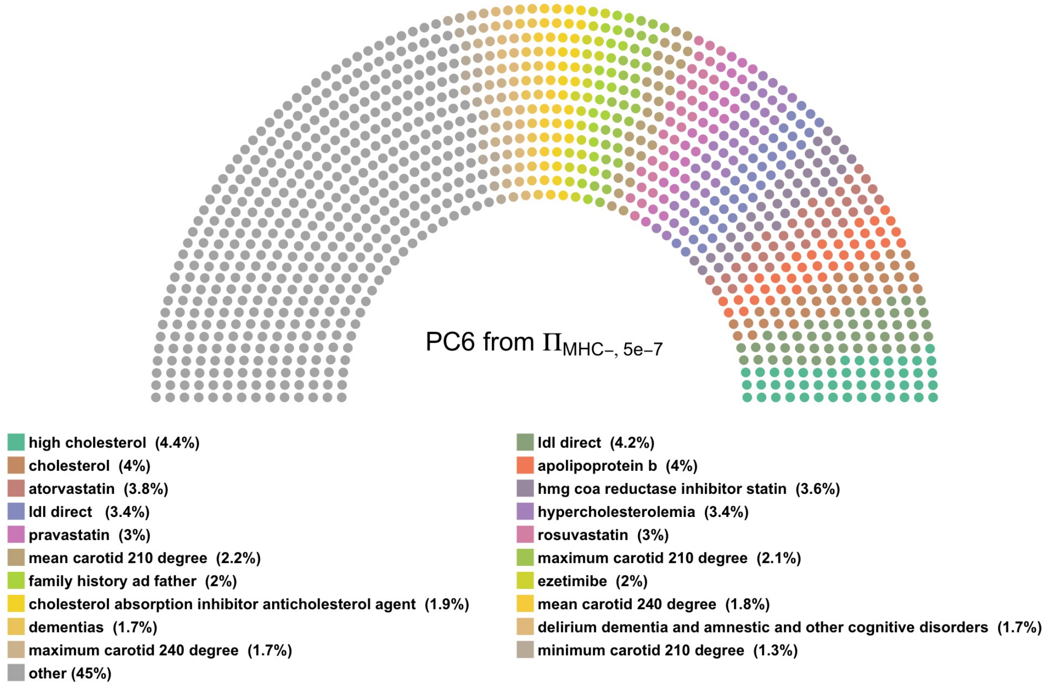

**((k))** This PC explained 1.82% of the total variation of the PRS matrix.

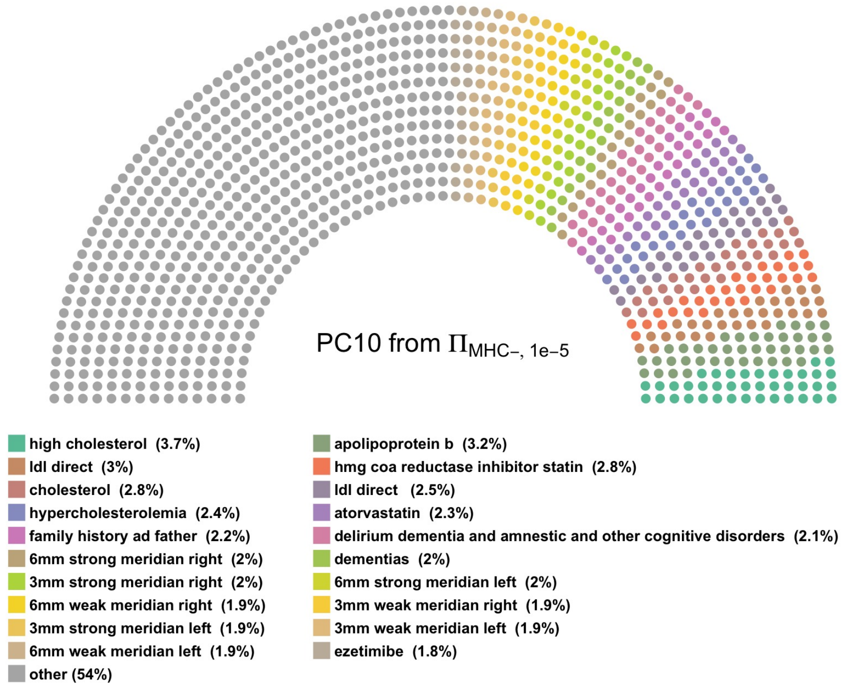

**((l))** This PC explained 0.39% of the total variation of the PRS matrix.

Fig. S11: Top 25 contributors, with their contribution, to PC6 and PC10, calculated from $\Pi_{MHC-,5e-07}$ and $\Pi_{MHC-,1e-05}$, are shown here. Each dot represents 0.4% of the PRS contribution to this PC.

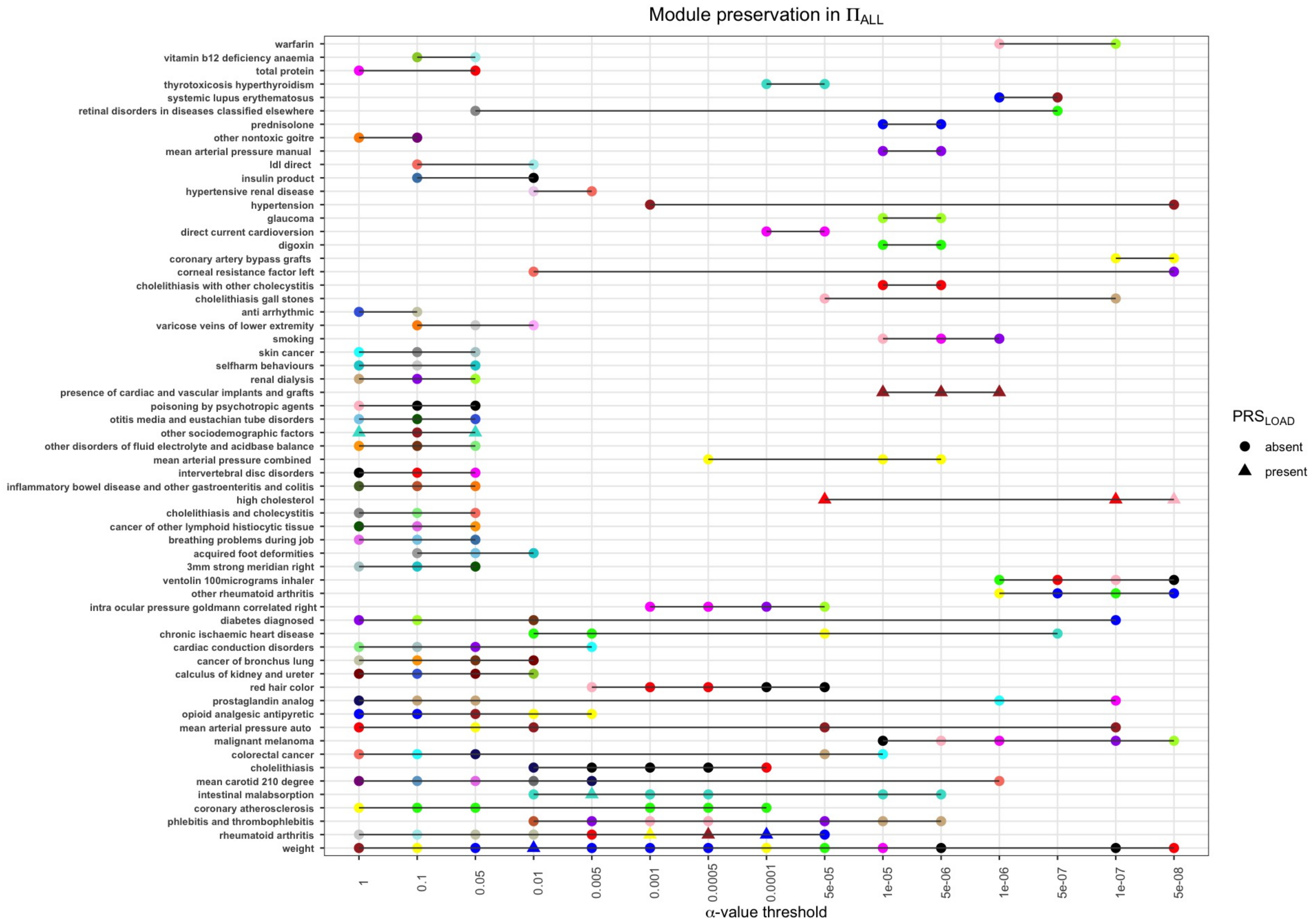

Fig. S12: Module preservation of the PRSs at every $\alpha$-value threshold for $\Pi_{ALL}$. On the y-axis, we present the most interconnected PRS in each module (shown as circles or triangles). The modules are outlined by their corresponding module name, and modules that include $PRS_{LOAD}$ are shown as triangles. We only included modules that are preserved at least two times across all 15 $\alpha$-value threshold. The rest of the modules are ordered from most conserved module (Weight) to least.

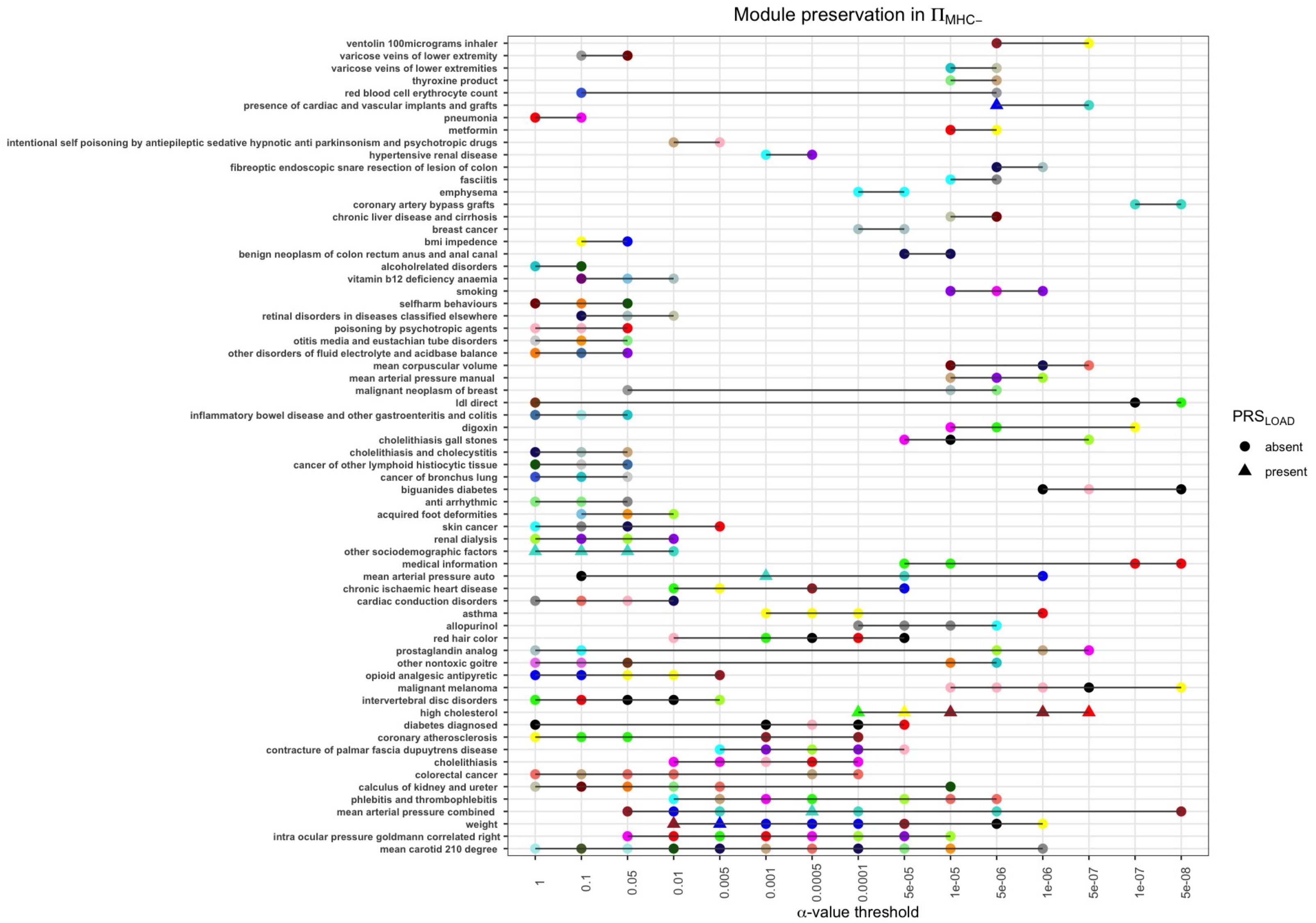

Fig. S13: Module preservation of the PRSs at every $\alpha$-value threshold for $\Pi_{MHC-}$. On the y-axis, we present the most interconnected PRS in each module (shown as circles or triangles). The modules are outlined by their corresponding module name, and modules that include $PRS_{LOAD}$ are shown as triangles. We only included modules that are preserved at least two times across all 15 $\alpha$-value threshold. The rest of the modules are ordered from most conserved module (mean carotid) to least.

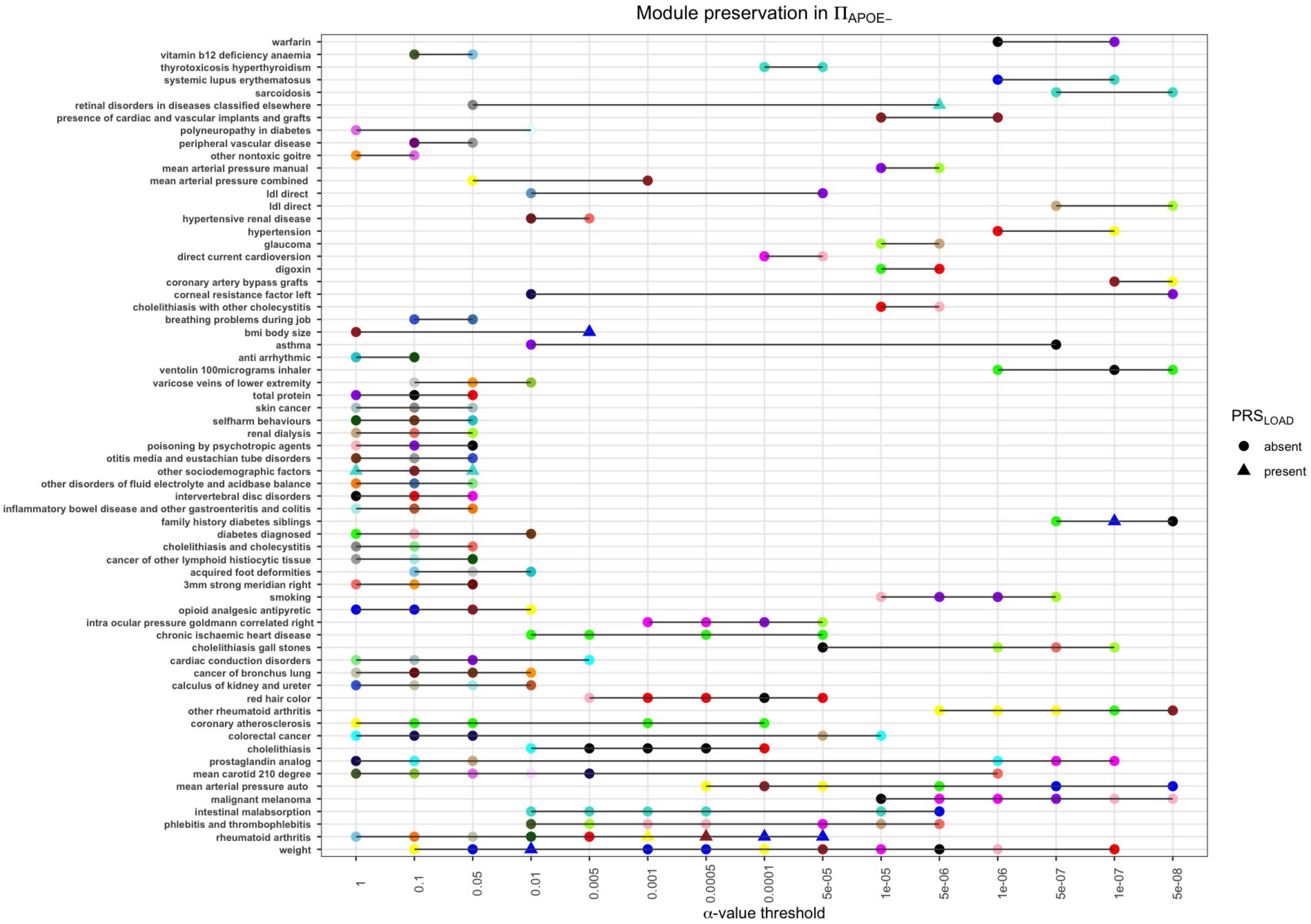

Fig. S14: Module preservation of the PRSs at every $\alpha$-value threshold for $\Pi_{APOE-}$. On the y-axis, we present the most interconnected PRS in each module (shown as circles or triangles). The modules are outlined by their corresponding module name, and modules that include $PRS_{LOAD}$ are shown as triangles. We only included modules that are preserved at least two times across all 15 $\alpha$-value threshold. The modules are ordered from most conserved module (Weight) to least. With the removal of the *APOE* region, we did not identify a module mostly interconnected with family history of AD dementia or any other AD-related PRS.

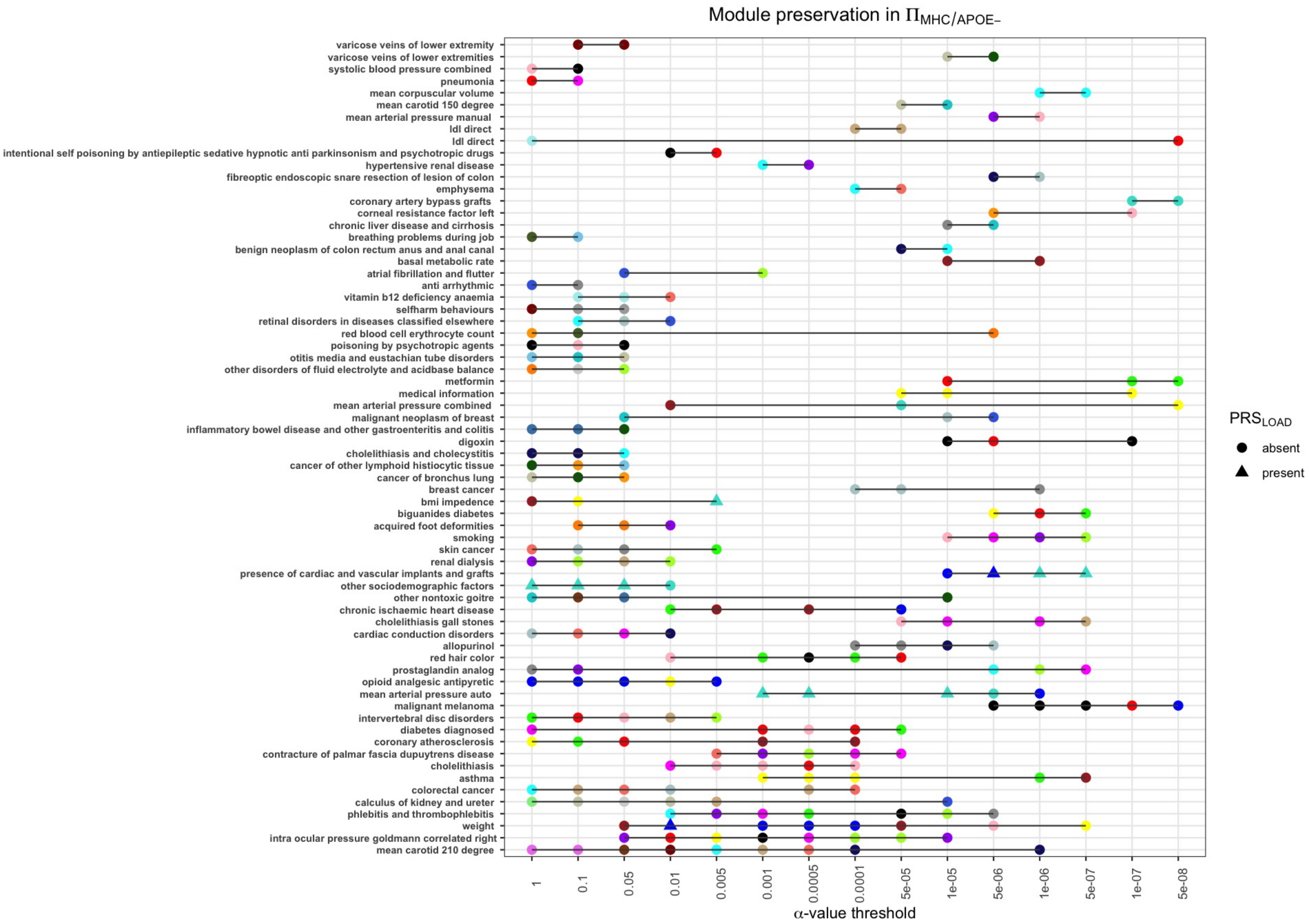

Fig. S15: Module preservation of the PRSs at every $\alpha$-value threshold for $\Pi_{MHC/APOE}$. On the y-axis, we present the most interconnected PRS in each module (shown as circles or triangles). The modules are outlined by their corresponding module name, and modules that include $PRS_{LOAD}$ are shown as triangles. We only included modules that are preserved at least two times across all 15 $\alpha$-value threshold. The modules are ordered from most conserved module (mean carotid) to least. With the removal of the *APOE* region, we did not identify a module mostly interconnected with family history of AD dementia or any other AD-related PRS.

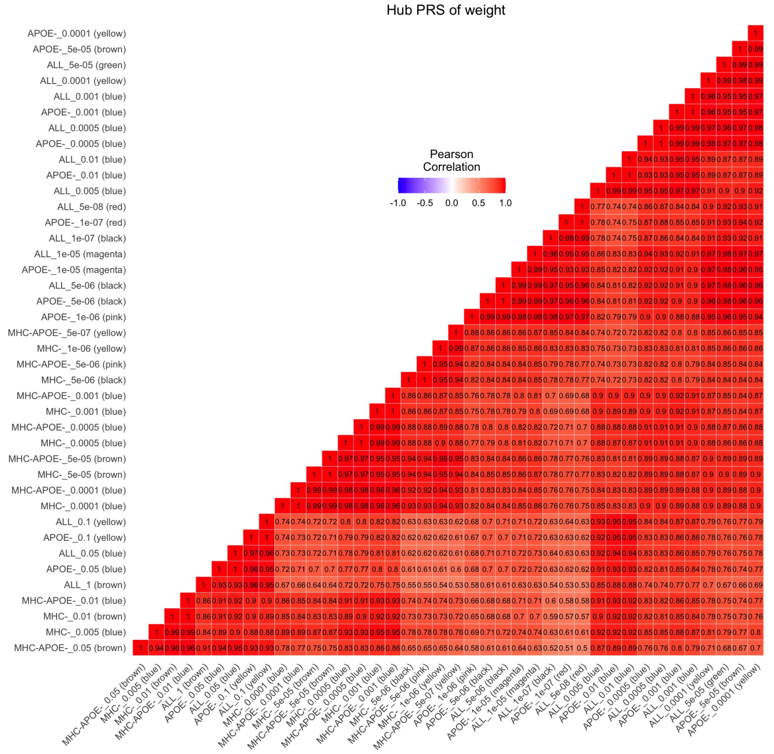

((m))

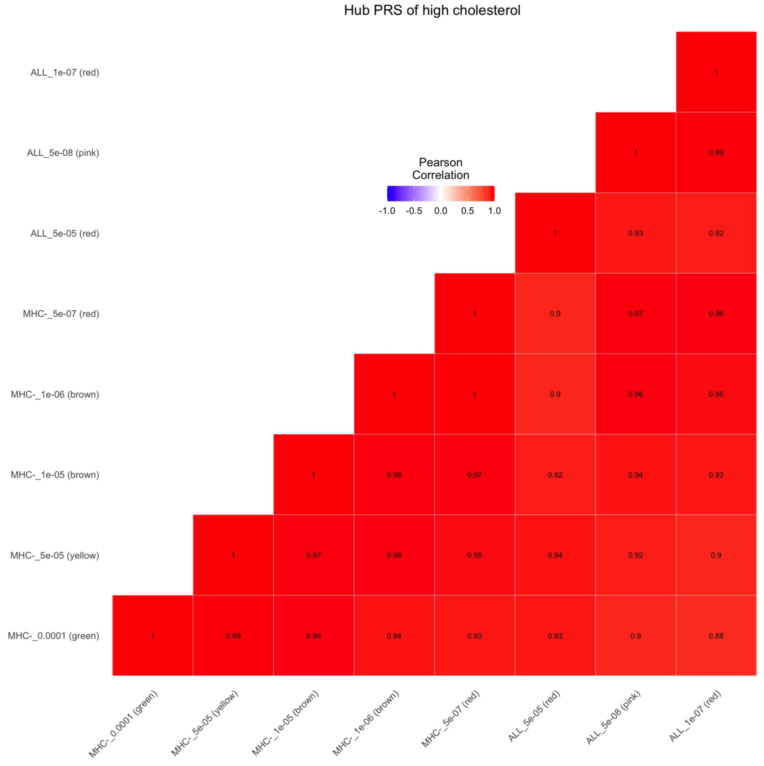

((n))

Fig. S16: Outlining the correlational structure of the hub PRSs for weight and high cholesterol across all $\alpha$-value thresholds for all 4 PRS matrices. We notice that in both heatmaps, the hub PRSs are highly correlated with each other, verifying the robustness of these modules.

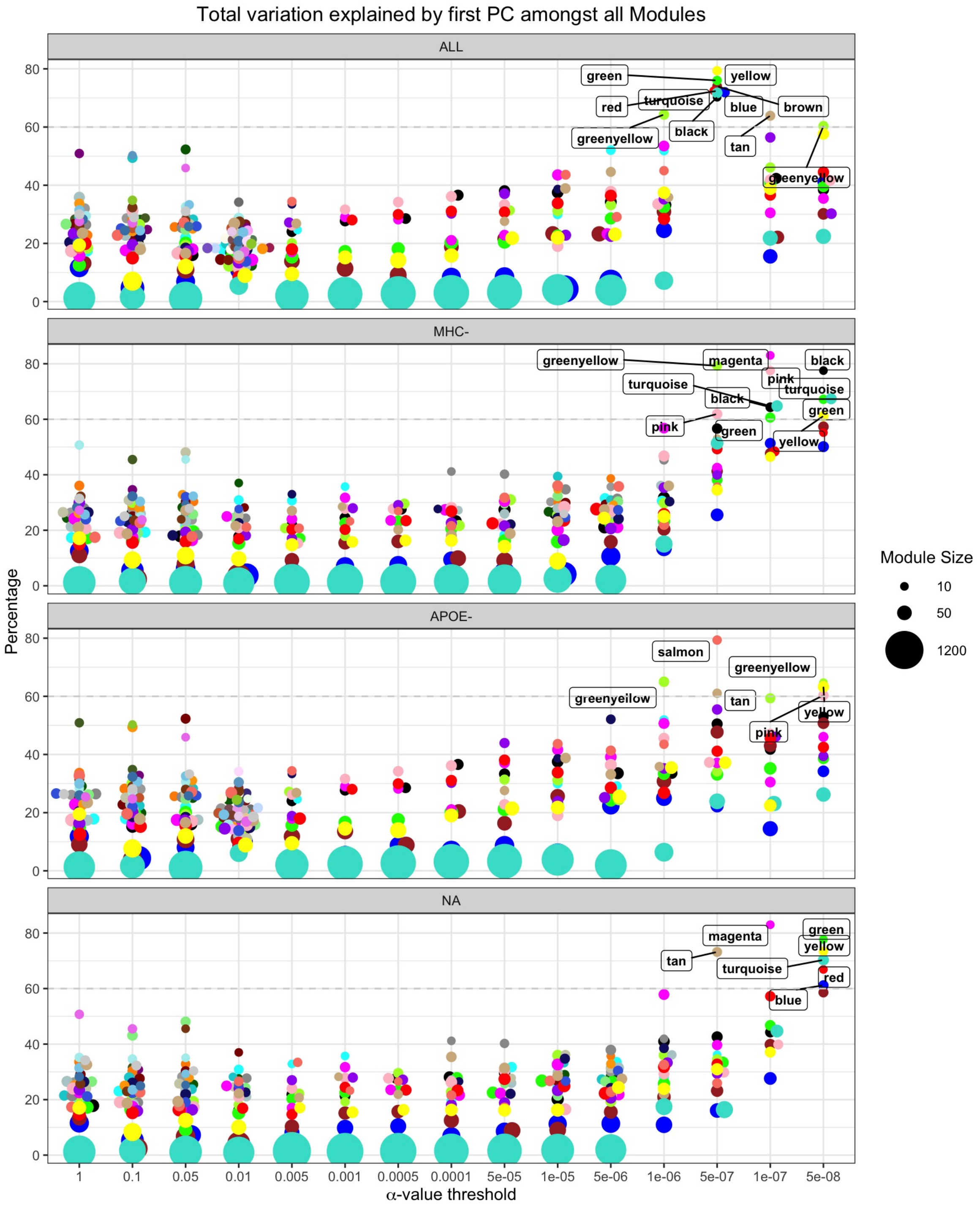

Fig. S17: This plot presents the total variation explained by PC1 in each module (generated by WGCNA). This measurement is highly dependent on the size of the module. The modules in the higher thresholds consist of a larger number of PRSs, explaining lower variance. Conversely, we can see smaller-sized modules at lower $\alpha$-value thresholds explaining higher variance (more than 40% on average). We identified modules, where their corresponding PC1 captured more than 60% of the total variance.

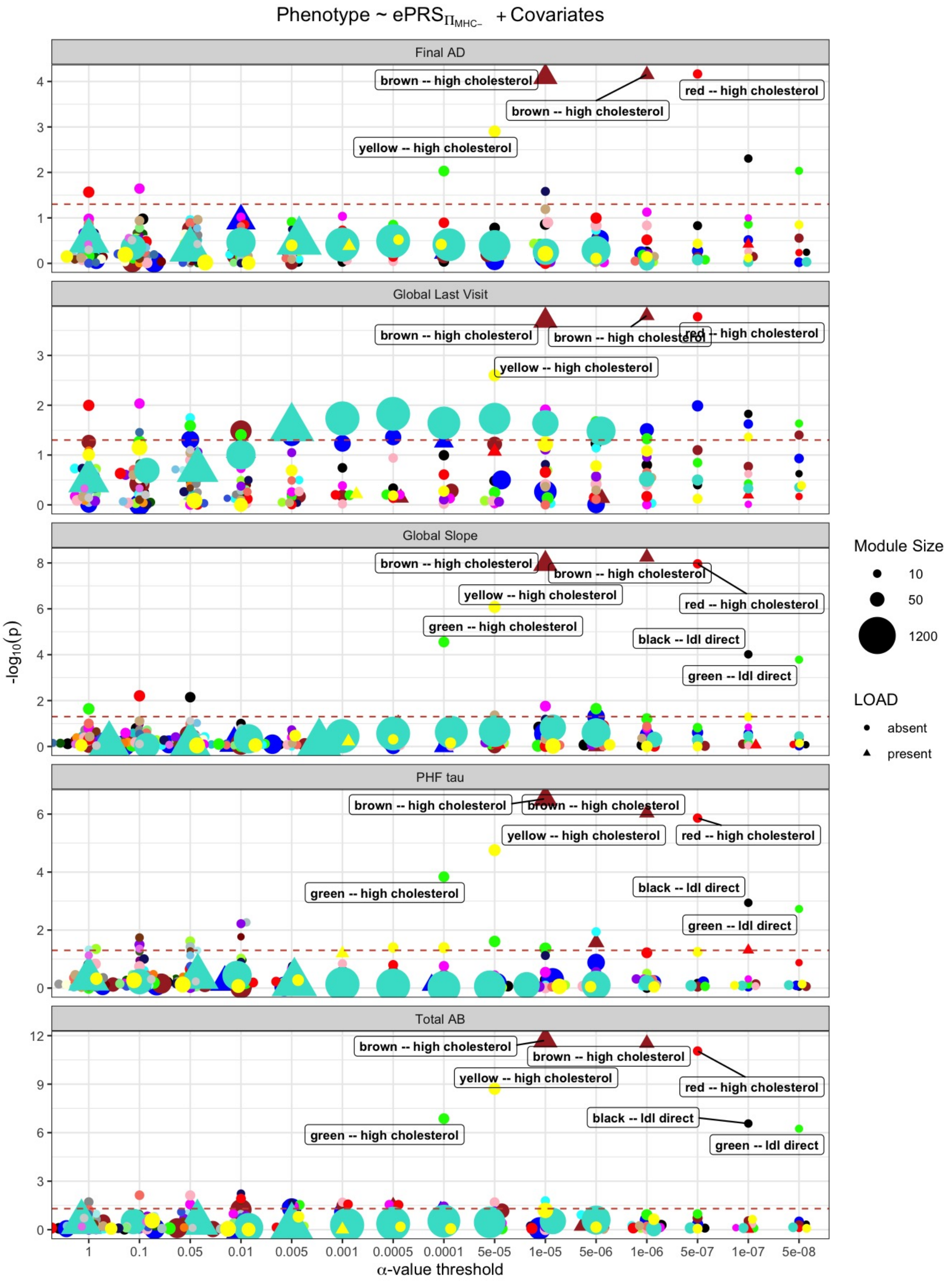

Fig. S18: Summarising the association of the aging phenotypes from the ROS/MAP study cohort and the ePRSs from $\Pi_{MHC-}$. The red dotted line represents 5% significance level. We outline the hub PRS of the highly associated module with the ageing phenotypes (q-value $\leq 0.05$). These modules were selected to test if they can improve the predictability ability of state-of-the-art $PRS_{LOAD}$.

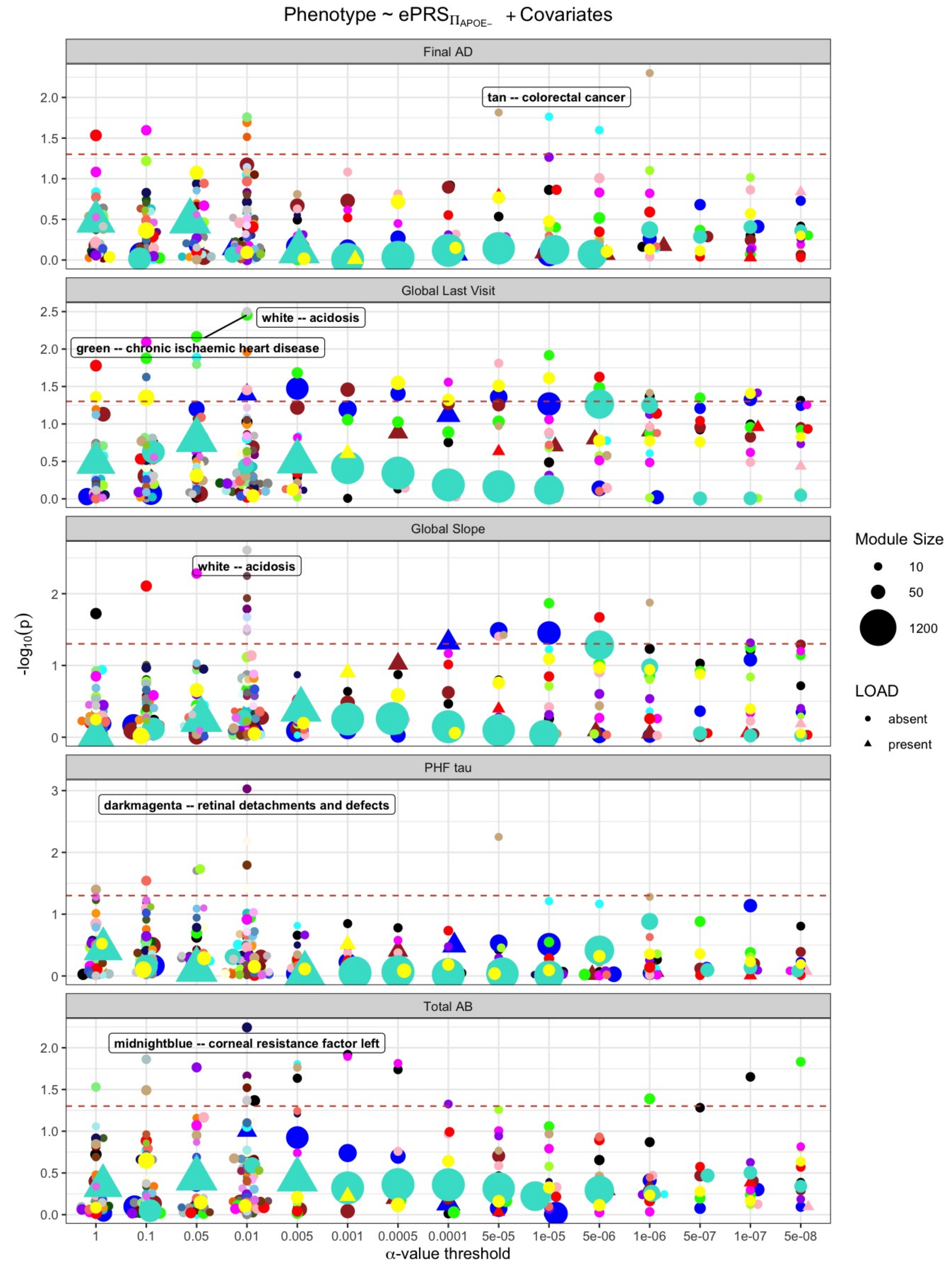

Fig. S19: Summarising the association of the aging phenotypes from the ROS/MAP study cohort and the ePRSs from $\Pi_{APOE-}$. The red dotted line represents 5% significance level. We outline the hub PRS of the highly associated module with the ageing phenotypes (smallest p-value, no module passed q-value $\leq 0.05$ testing). These modules were selected to test if they can improve the predictability ability of state-of-the-art $PRS_{LOAD}$.

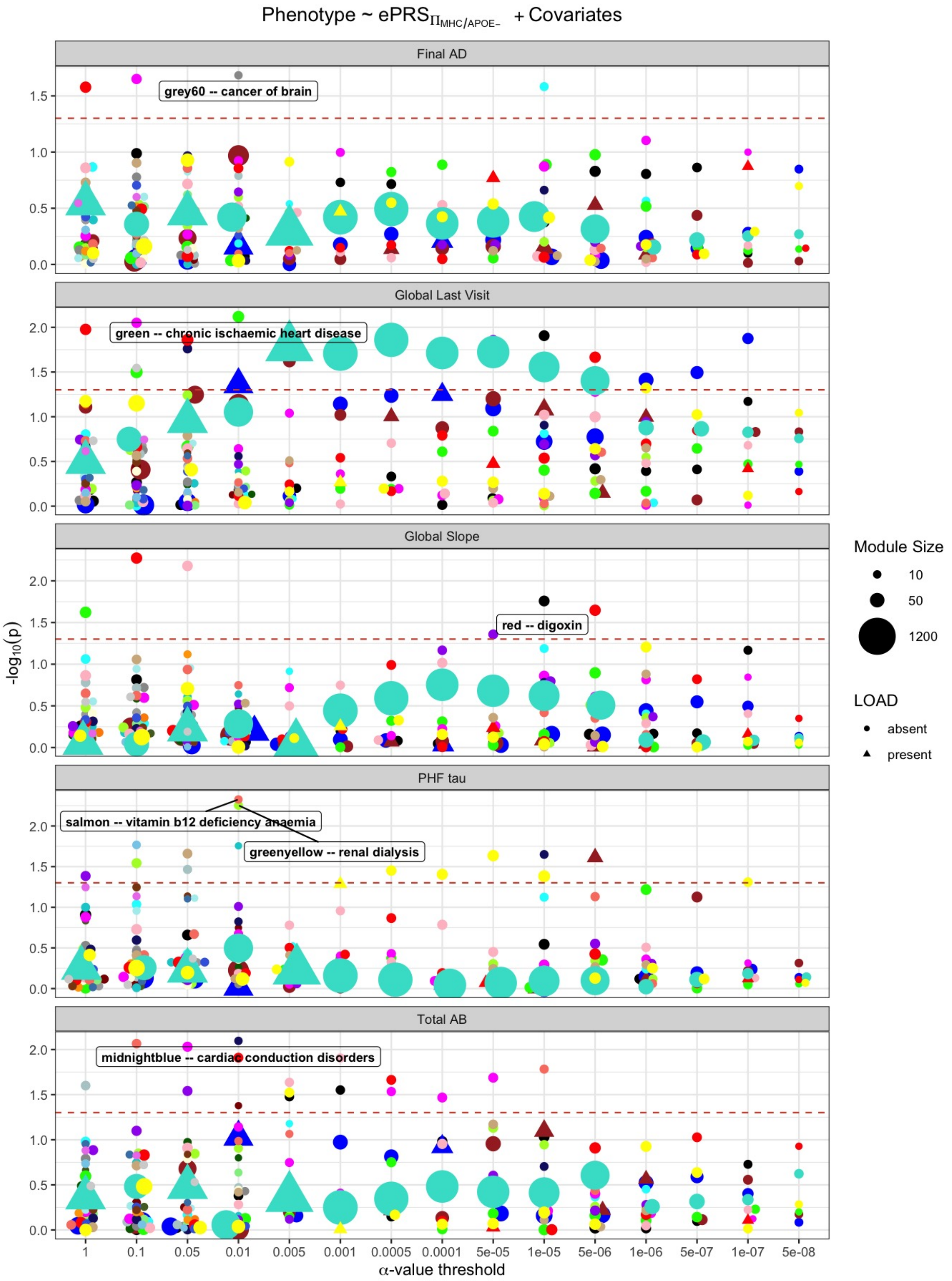

Fig. S20: Summarising the association of the aging phenotypes from the ROS/MAP study cohort and the ePRSs from $\Pi_{MHC/APOE-}$. The red dotted line represents 5% significance level. We outline the hub PRS of the highly associated module with the ageing phenotypes (smallest p-value, no module passed q-value $\leq$ 0.05 testing). These modules were selected to test if they can improve the predictability ability of state-of-the-art $PRS_{LOAD}$.

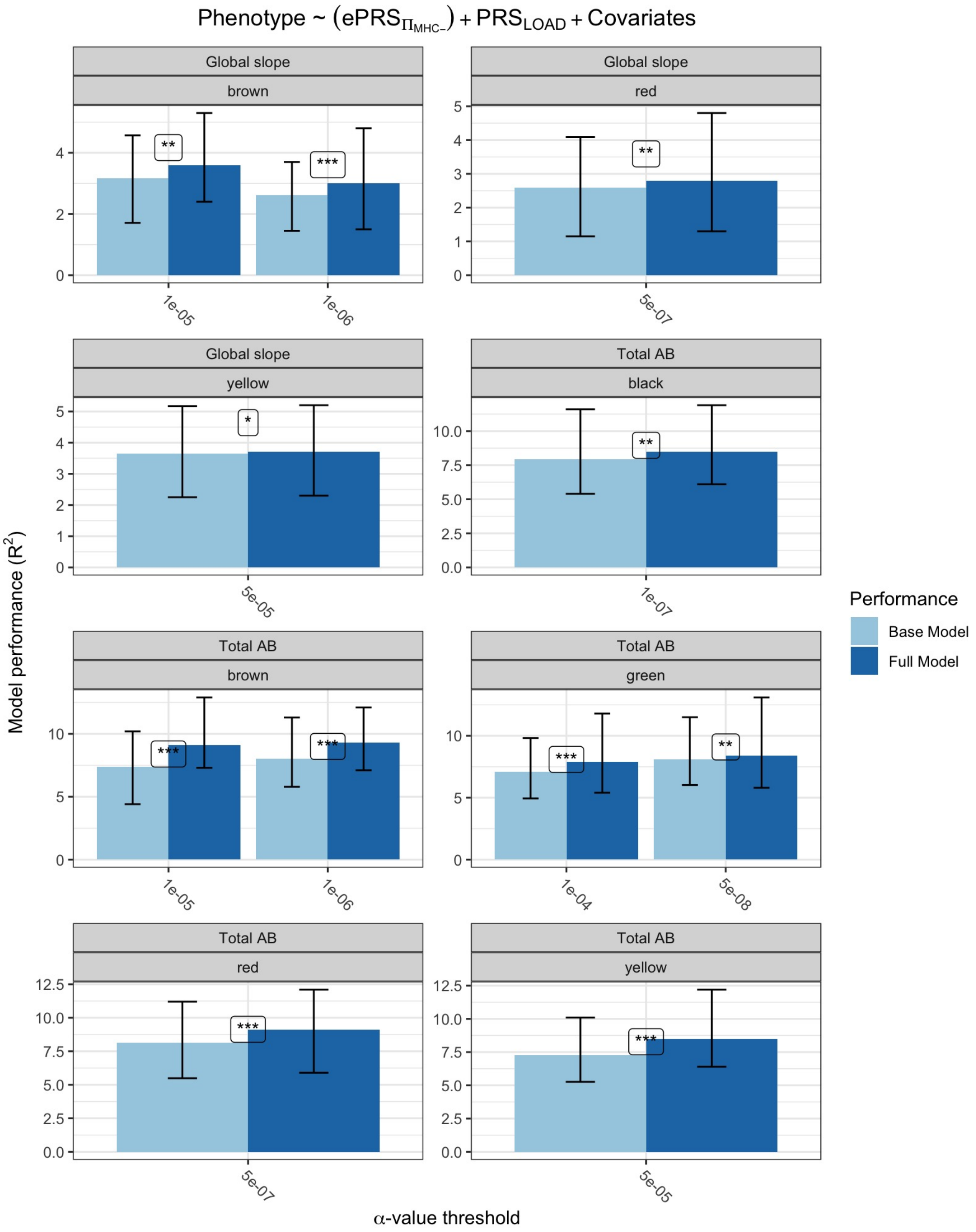

Fig. S21: Summarizing the results of the multivariate association of modules from $\Pi_{MHC-}$. False discovery rate (FDR) correction (i.e. $q$-value [3]) was used to mitigate multiple testing concerns among all ePRSs (modules) on the likelihood ratio test between the full and base models. We selected ePRSs (modules) with q-value $\leq 0.05$ and report it here. For continuous response variables (Beta Amyloid ($\beta$-amyloid) protein, PHF Tau, cognitive global random slope, cognitive global random slope at last visit) we report the .632 bootstrapped variation explained ($R^2$) by the covariates for the base (model 2) and full model (model 3). Additionally, we indicate the p-value of the likelihood ratio test between the full and base models. *$p \leq 0.05$, **$p \leq << 0.01$, ***$p \leq 0.001$. All p-values are reported at an uncorrected level. Note that the confidence intervals were measured out of the bootstrapping loop.
